# Dual-mode action of scalable, high-quality engineered stem cell-derived SIRPα-extracellular vesicles for treating acute liver failure

**DOI:** 10.1101/2024.05.24.592430

**Authors:** Seohyun Kim, Yoon Kyoung Kim, Seonghyun Kim, Yong-Soon Choi, Inkyu Lee, Hyemin Joo, Jaehyun Kim, Minjeong Kwon, Seryoung Park, Min Kyoung Jo, Yoonjeong Choi, Theresa D’Souza, Jae Woong Jung, Elie Zakhem, Stephen Lenzini, Jiwan Woo, Hongyoon Choi, Jeongbin Park, Seung-Yoon Park, Gi Beom Kim, Gi-Hoon Nam, In-San Kim

## Abstract

Acute liver failure (ALF) is a critical inflammatory condition characterized by rapid hepatocyte death, impaired liver regeneration due to the delayed removal of necroptotic cells, and high mortality rates. This study introduces a novel dual-mode action therapeutic approach using extracellular vesicles expressing Signal Regulatory Protein Alpha (SIRP-EVs) derived from genetically engineered mesenchymal stem cells (MSCs). These SIRP-EVs are designed to concurrently resolve necroptosis and promote liver regeneration. Our studies identified CD47 and SIRPα as promising therapeutic targets for ALF. We developed a scalable 3D bioreactor-based process that produces high-purity SIRP-EVs, which preserve MSC properties and achieve significant production levels. SIRP-EVs target both macrophages and necroptotic hepatocytes in ALF models, enhancing macrophage phagocytic activity against necroptotic cells via CD47 blockade and promoting liver regeneration by reprogramming macrophages with MSC-derived cargo. Comprehensive *in vitro* and *in vivo* studies demonstrate that SIRP-EVs decrease CD47^+^ necroptotic cells and promote liver regeneration in ALF models, leading to reduced liver damage markers and enhanced survival rates. These findings highlight the potential of SIRP-EVs as a dual-mode action therapeutic for ALF, offering promising prospects for their application in other inflammatory diseases. Moreover, these results pave the way for advancing engineered EV-based therapies toward clinical implementation.

## Introduction

Acute inflammation, whether caused by infectious or non-infectious origins, often leads to severe outcomes, including organ failure^1^. Acute liver failure (ALF) exemplifies this issue, with a mortality rate exceeding 50 %, largely due to limited therapeutic options^2^. ALF is characterized by the rapid death of hepatocytes, exacerbated by impaired clearance of these cells and insufficient regenerative responses, underscoring the urgent need for effective therapeutic strategies.

Recent studies highlight necroptosis, a form of regulated necrosis, as a significant contributor to ALF^3–5^. Triggered by inflammatory cytokines, necroptosis compromises cell membranes, releasing intracellular contents that further exacerbate inflammation^6^. While macrophages typically engulf dying cells via efferocytosis, necroptotic cells often evade this clearance by expressing high levels of CD47-a ‘don’t eat me’ signal-that prevents macrophage-mediated clearance^7–9^. Furthermore, inflammation triggered by uncleared necroptotic cells can polarize highly plastic macrophages towards a pro-inflammatory phenotype, perpetuating liver injury^10,11^. This barrier to resolution complicates inflammation control and hinders liver regeneration, underscoring the therapeutic challenges in managing ALF.

To address these unmet medical needs, we developed a dual-mode action therapeutic approach using extracellular vesicles expressing SIRPα, the natural ligand of CD47, derived from engineered human mesenchymal stem cells (SIRP-EVs). Initial analyses of ALF models revealed a direct correlation between necroptosis and CD47 expression, supported by comprehensive protein and transcriptomic analyses. We also established a rigorous SIRP-EV manufacturing process that ensures scalability, high purity, and yield, while retaining the inherent properties of MSCs. Additionally, we explored the ability of SIRP-EVs to target both macrophages and necroptotic hepatocytes in ALF models. Our strategy effectively enhances macrophage functionality, specifically increasing phagocytic activity against necroptotic hepatocytes via CD47 blockade and promoting liver regeneration through the reprogramming of macrophages with MSC-derived cargo. This study demonstrates the therapeutic potential of SIRP-EVs in treating ALF, marking a significant advancement in addressing this critical healthcare challenge.

## Results

### Elevated CD47 expression in necroptotic hepatocytes hallmarks ALF

We developed various animal models of ALF to understand complexity, each model induced by different etiological factors. These models employ pharmacological agents such as Acetaminophen (APAP), Thioacetamide (TAA), Carbon tetrachloride (CCl4), and a combination of Lipopolysaccharide (LPS) with D-galactosamine (D-galN), each representing distinct pathways of hepatic injury. Specifically, APAP-induced hepatotoxicity results from its metabolic conversion into toxic metabolites. TAA and CCl4 induce liver damage through bioactivation to reactive intermediates causing hepatocellular damage. Meanwhile, the LPS/D-galN model mimics pathogen-induced liver failure^12,13^. These models provide valuable insights into the pathophysiological mechanisms of ALF and assist in identifying novel therapeutic targets for various liver injury etiologies.

Our analysis showed significant increases in serum aspartate aminotransferase (AST) and alanine aminotransferase (ALT) levels across all ALF models, indicative of severe hepatic injury **(Fig. 1A and Supplementary Fig. 1)**. Hematoxylin & Eosin (H&E) staining and immunohistochemistry (IHC) on liver tissues revealed extensive cellular death and a marked increase in CD47 expression in all ALF models compared to normal liver tissue **(Fig. 1A and Supplementary Fig. 1)**. Subsequent studies aimed to identify liver cell populations with elevated CD47 expression following ALF induction. Flow cytometry analysis revealed a significant upregulation of CD47 in hepatocytes from ALF conditions compared to normal **(Fig. 1B and Supplementary Fig. 2A)**. This increase was further confirmed by western blot analysis of hepatocytes isolated from ALF-affected liver tissues, showing a notable enhancement in CD47 expression **(Fig. 1C)**. Annexin V/7-AAD assays indicated a significant rise in the late apoptotic/necroptotic cell population in CD47-high hepatocytes, especially in the APAP-ALF model **(Fig. 1D and Supplementary Fig. 2B)**. This was associated with overexpression of necroptotic markers, such as RIP3 and pMLKL, in ALF tissues **(Fig. 1E and Supplementary Fig. 2C, D)**. Confocal microscopy analysis corroborated a direct correlation between CD47 expression and the pMLKL in ALF-affected liver tissues **(Fig. 1F)**. These findings suggest a significant upregulation of CD47 expression in necroptotic hepatocytes across ALF models compared to normal liver.

**Fig. 1:**
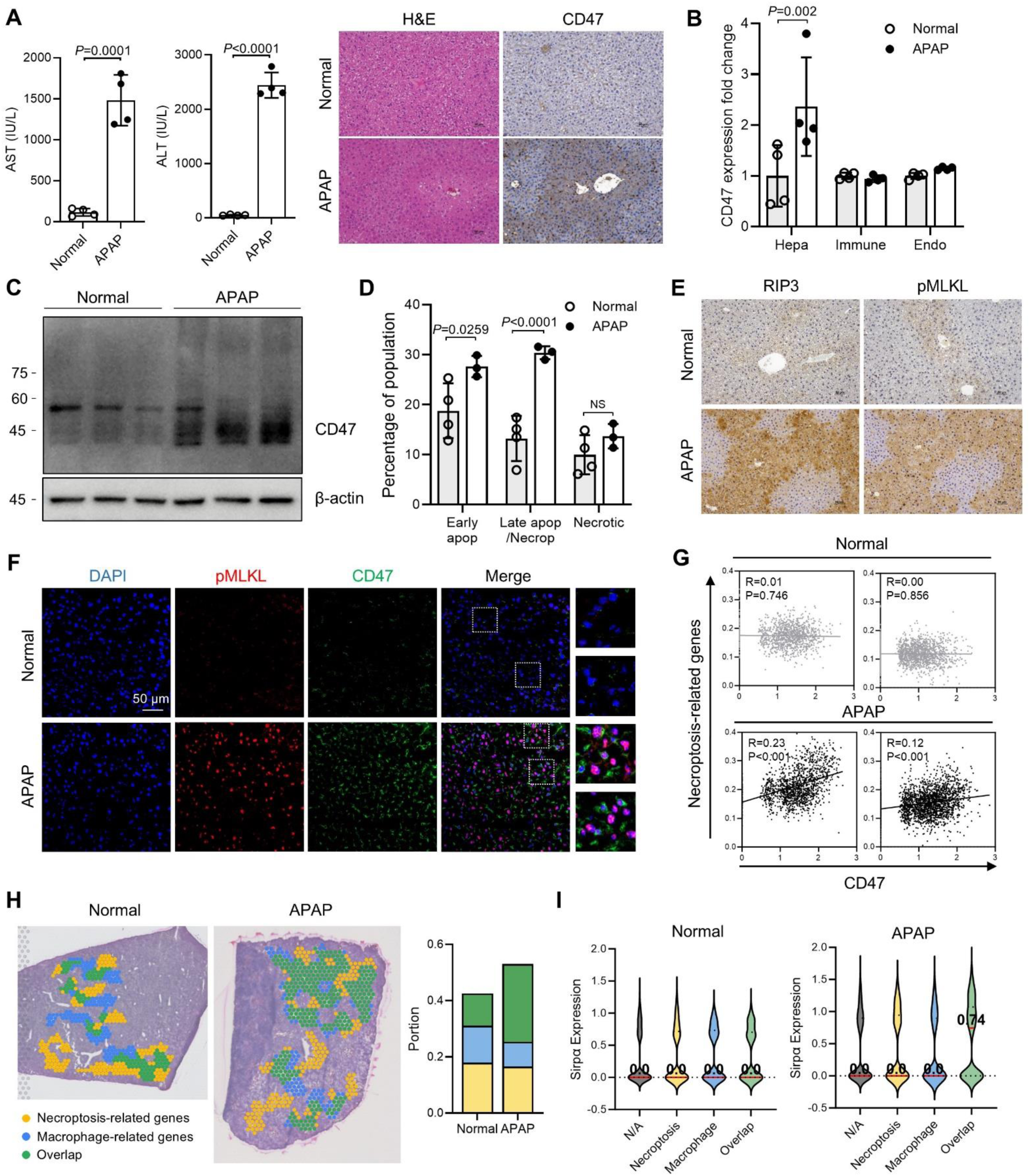
CD47 is overexpressed on necroptotic hepatocytes in the damaged liver of ALF models. (A) Biochemical evaluation (AST and ALT levels) of the ALF model after APAP induction (left). Representative images of H&E and CD47 staining of liver samples from an ALF model induced by APAP (right) (n=4). (B) CD47 expression levels of cell populations within liver tissue (n=4). (C) CD47 expression in hepatocytes from normal and APAP-ALF livers (n=3). (D) Graphical representation of early apoptotic (Annexin V^+^/7-AAD^-^), late apoptotic/necroptotic (Annexin V^+^/7-AAD^+^), and necrotic (Annexin V^-^/7-AAD^+^) cell populations in liver hepatocytes from normal and APAP-ALF mice (n=3 or 4 per group). (E) Expression of RIP3 and pMLKL in liver samples from normal and APAP-ALF groups. (F) Representative confocal images of liver sections from normal and APAP-ALF mice. Scale bar, 50 μm. (G) Scatter plot visualization of the Spearman correlation (R) between 121 necroptosis gene scoring and RNA levels of CD47 across samples. (H) STopover analysis to map the spatial overlap and interactions between cell types in liver tissue from normal and APAP-ALF mice. Highlights include 121 necroptosis gene scores (yellow) and macrophage RNA levels (blue), with overlapping areas shown in green. The plots below represent the three regions. (I) Violin plot visualizing SIRPα expression levels in different regions based on spatial correlation results. The numbers marked on the plot represent the median values of the SIRPα expression. Bar graph data are presented as mean ± S.D. Statistical significance was determined by two-tailed unpaired student’s t-test (A) and two-way ANOVA with Sidak’s post-hoc test (B and D). Hepa, Hepatocytes; Immune, Immune cells; Endo, Endothelial cells.

Spatial transcriptomic (ST) analysis of liver tissue from the APAP-induced ALF model (APAP- ALF) demonstrated a distinct spatial association between necroptosis-related genes and CD47 expression, differentiating ALF liver tissues from normal tissues **(Fig. 1G)**. In contrast, apoptosis-related genes showed little to no evident correlation with CD47 expression (**Supplementary Fig. 3A**). This analysis was conducted using a publicly available ST dataset (GEO accession number GSE223560), processed through the Seurat pipeline to focus on spatial features, including necroptosis and CD47 expression, with correlations assessed via Spearman analysis.

Using the CellDART algorithm^14^ cell types were mapped by integrating ST data with reference single-cell RNA-seq data, and STopover^15^ was employed to examine the colocalization of necroptosis- and macrophage-related gene expression spots. This analysis was expanded to identify spatial overlap between necroptosis and macrophage-related genes, which was more pronounced in ALF liver tissues than in healthy liver (**Fig. 1H**). However, there was no significant spatial overlap observed between apoptosis-related genes and macrophage-related genes in ALF liver compared to normal liver (**Supplementary Fig. 3B**). Additionally, there was a significant increase in SIRPα expression at the colocalized sites of these genes **(Fig. 1I)**. Previous studies have indicated that macrophages expressing high levels of SIRPα tend to be located around cells that overexpress CD47^9,16^. Therefore, these results suggest that the CD47/SIRPα interaction, which inhibits efferocytosis in necroptotic hepatocytes, is upregulated in ALF and could serve as a therapeutic target for ALF.

### 3D bioreactor-based scalable purification process produces SIRP-EVs with high purity and preserves MSC characteristics

To address inflammatory diseases associated with upregulated CD47, we genetically engineered human bone marrow-derived MSCs to express SIRPα creating SIRPα-modified MSCs (SIRP-MSCs) through stable transduction with lentiviral vectors encoding the SIRPα. These cells retained the hallmark differentiation capabilities of naive MSCs, including differentiation into adipocytes, osteoblasts, and chondrocytes **(Fig. 2A)**. Importantly, SIRP- MSCs maintained the expression of key MSC markers (CD73, CD90, CD105, and CD166) and did not express hematopoietic and endothelial markers (CD14, CD34, and CD45), as confirmed by flow cytometry **(Fig. 2B)**. Multiplex ELISA analysis showed that the angiogenic cytokine secretion profiles of SIRP-MSCs were similar to those of non-modified MSCs (C-MSCs), indicating that the genetic modification did not alter the inherent properties of MSCs **(Fig. 2C)**. Analysis of inter- and intra-donor variability showed consistent proliferation rates, EV yield, and the proportion of SIRPα-positive cells across biological replicates, demonstrating a high level of reproducibility in the modification process **(Fig. 2D)**.

**Fig. 2:**
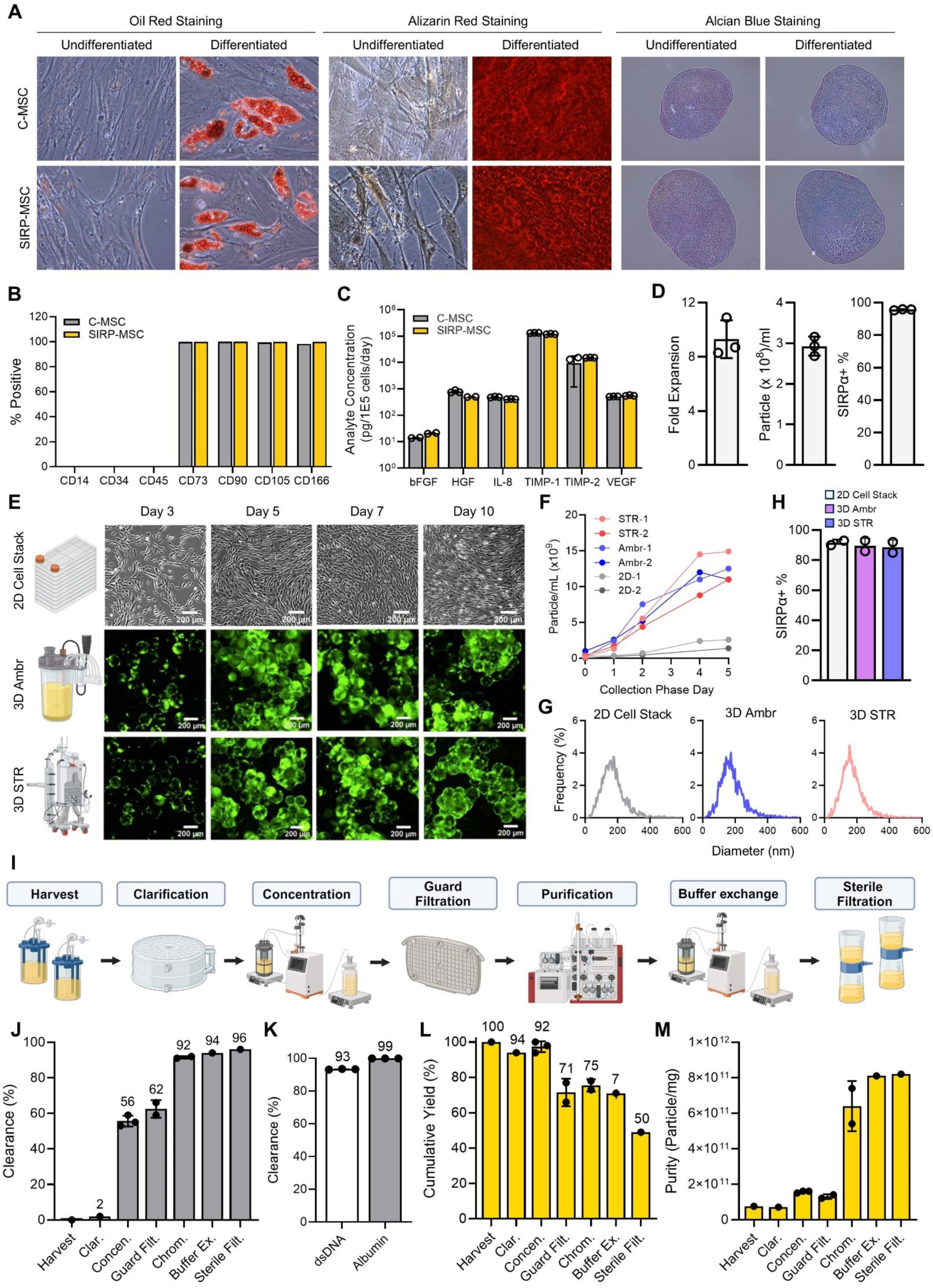
Scalable SIRP-EV production from engineered MSCs using 3D Bioreactor systems, ensuring high purity and integrity. (A) Trilineage differentiation of engineered MSCs: adipogenic differentiation by lipid droplets (left), osteogenic differentiation by mineralized matrices (middle), and chondrogenic differentiation by cartilage matrix (right). (B) Surface marker expression on SIRP-MSCs post-transduction and control C-MSCs, confirming MSC phenotype with markers CD73, CD90, CD105, CD166, and absence of CD14, CD34, CD45. (C) Cytokine secretion profiles (FGF, HGF, IL-8, TIMP-1, TIMP-2, VEGF) of C-MSCs and SIRP-MSCs (n=2 or 3). (D) Inter- and intra-donor variability in SIRP-MSCs shown by fold expansion 5 days post-thaw (left), nanoparticle concentration at final harvest (middle), and percentage of SIRPα-stained cells (right). Two independent vials from a single donor and one from a second donor were evaluated (n=3). (E) Visualization of cells in 2D CellSTACKs and 3D bioreactor systems (Ambr250 and STR) (F) The time course measurements of particle concentration during two independent runs for each of the 2D CellSTACKs (2D), 3D Ambr250 (Ambr), and 3D STR (STR) systems. (G) Size distributions of particles using NTA from 2D CellSTACKs and 3D bioreactor systems (Ambr250 and STR). (H) SIRPα expression levels of MSCs before harvest from STR runs, with controls from Ambr and 2D systems (n=2). (I) Schematic of the downstream processing flow for SIRP-EV isolation, encompassing clarification to sterile filtration. (J-L) In-process analytics results for SIRP-EV: (J) protein, (K) dsDNA, and human serum albumin clearance, and (L) cumulative yield of SIRP-EV. (M) The purity of SIRP-EV per step is quantified as the particle count per mg protein. (B-D, H, J-M) Bar graph data are presented as mean ± S.D. (J, L, M) Bars represent the mean of independent process replicate points using mean analytical test results. (K) dsDNA and albumin tests included n=3 technical test replicates. The numbers displayed on a bar graph represent the exact numerical values of the bar. Clar, clarification; Concen, concentration; Guard Filt, guard filtration; Chrom, chromatography; Buffer Ex, buffer exchange; Sterile Filt; sterile filtration

For the efficient isolation of SIRP-EVs from SIRP-MSCs, we employed a 3D bioreactor system utilizing microcarriers in the upstream process. The efficacy of this system was assessed by comparing the growth of SIRP-MSCs in both small-scale (250ml) Ambr250 (Ambr) and large-scale (15L) Biostat STR^®^ (STR) bioreactors with traditional 2D CellSTACKs as the control. Daily inspections of the microcarrier suspensions in the bioreactor vessels (STR and Ambr) confirmed a consistent suspension of cells and microcarriers, ensuring an evenly mixed culture **(Supplementary Fig. 4A)**. Critical process parameters such as temperature, dissolved oxygen, and agitation rates, were meticulously controlled. The metabolite profiles from the bioreactor vessels were closely aligned, indicating consistent process conditions **(Supplementary Fig. 4B)**. Cells were monitored using a live stain, demonstrating clear attachment and expansion on microcarriers over a 10-day culture period **(Fig. 2E)**. Daily measurements of particle count from the culture media, following the replacement of growth media with EV collection media, showed that 3D bioreactors produced significantly more EVs than 2D cultures **(Fig. 2F, G and Supplementary Fig. 4C)**. Additionally, the expression of SIRPα in MSCs remained stable throughout the collection period, up to the last day **(Fig. 2H)**.

The scalability of our upstream process development was validated through experiments conducted in a 50 L STR bioreactor, with a 250 ml Ambr system serving as the scale-down. Comprehensive analyses including macroscopic and microscopic imaging, metabolite profiles, cell proliferation, cell viability, and EV counts per ml demonstrated consistency across scales, confirming a successful 200-fold scale-up in a 3D bioreactor **(Supplementary Fig. 5)**. The downstream process comprised depth filter filtration for clarification, tangential flow filtration for concentration, chromatography for purification, followed by an additional tangential flow filtration for buffer exchange and further concentration, and concluding with sterile filtration prior to vialing **(Fig. 2I)**. The final SIRP-EVs showed a significant cumulative reduction in double-stranded DNA (dsDNA) and albumin by 93% and 99%, respectively, with a 96% clearance of total protein content from the initial culture medium **(Fig. 2J, K and Supplementary Fig. 6A-C)**. After the downstream process, the purity of the isolated SIRP- EVs was remarkably high, reaching 8 x 10^11^ particles per mg of proteins, with a cumulative yield, or particle recovery rate, of 50% **(Fig. 2L, M and Supplementary Fig. 6D)**. These results demonstrate that our process development for the 3D-based EV isolation can efficiently separate SIRP-EVs with high purity and yield.

### SIRP-EVs carry MSC-derived protein cargos and present a safe CD47 blockade

Naïve MSC-derived EVs (C-EVs) are recognized for their effectiveness in treating inflammatory diseases through the transfer of miRNA and protein cargos^17,18^. To determine if SIRP-EVs replicate a similar MSC-derived cargo profile, we produced C-EVs using the same upstream and downstream processes, achieving high particle recovery rates and purity **(Supplementary Fig. 7)**. Initial analysis of the RNA content using RNA electrophoresis revealed negligible RNA levels in SIRP-EVs **(Supplementary Fig. 8A)**. To evaluate the potential impact of processing methods on RNA cargo and to perform more precise RNA quantification, we compared RNA quantities using the RiboGreen assay. This comparison was made between EVs isolated through research-grade processes—including serial centrifugation, tangential flow filtration, and ultracentrifugation—and those in C-EVs and SIRP-EVs isolated using our established 3D bioreactor-based process. The results indicated that EVs isolated through research-grade processes contained more RNA than those isolated using the 3D-based system **(Supplementary Fig. 8B)**. These findings suggest that RNA content of EVs can vary significantly depending on the isolation and purification process used.

We utilized liquid chromatography with tandem mass spectrometry (LC-MS/MS) to analyze the protein profiles of SIRP-EVs, identifying 7,391 proteins. This confirmed that the protein cargos of SIRP-EVs closely mirror those of C-EVs **(Fig. 3A, B)**, while also demonstrating overexpression of SIRPα in SIRP-EVs **(Fig. 3C)**. Further experiments were conducted to verify the EV nature of SIRP-EVs. Western blot analysis confirmed the presence of EV markers, including CD81, CD9, CD63, and TSG101, as well as the engineered expression of SIRPα, while confirming the absence of the non-EV marker, prohibitin^19^ **(Fig. 3D and Supplementary Fig. 9)**. Cryo-transmission electron microscopy (cryo-TEM) revealed that SIRP-EVs, like C- EVs, are spherical nanoparticles characterized by a lipid bilayer **(Fig. 3E and Supplementary Fig. 10)**. Dynamic light scattering (DLS) assays showed that SIRP-EVs, similar to C-EVs, exhibit a uniform particle size distribution with average diameters of approximately 200 nm, indicative of no particle aggregation and a high degree of monodispersity, as reflected by a polydispersity index close to 0.2 (SIRP-EV: 0.190, C-EV: 0.187) **(Fig. 3F)**. Zeta potential (ZP) measurements recorded SIRP-EVs and C-EVs at −22.56 mV and −22.30 mV by nanoparticle tracking analysis (NTA), respectively, with both demonstrating over 95% lipidation, confirmed through MemGlow staining **(Fig. 3G)**. Further analysis of the over-represented Gene Ontology Biological Processes (GOBPs) within the SIRP-EV protein cargo revealed an enrichment of adhesion and binding proteins that promote directed and prolonged binding of C-EVs at sites of inflammation^20^ **(Fig. 3H)**. Collectively, these findings demonstrate that SIRP-EVs maintain the fundamental characteristics of EVs, comparable to those of C-EVs.

**Fig. 3:**
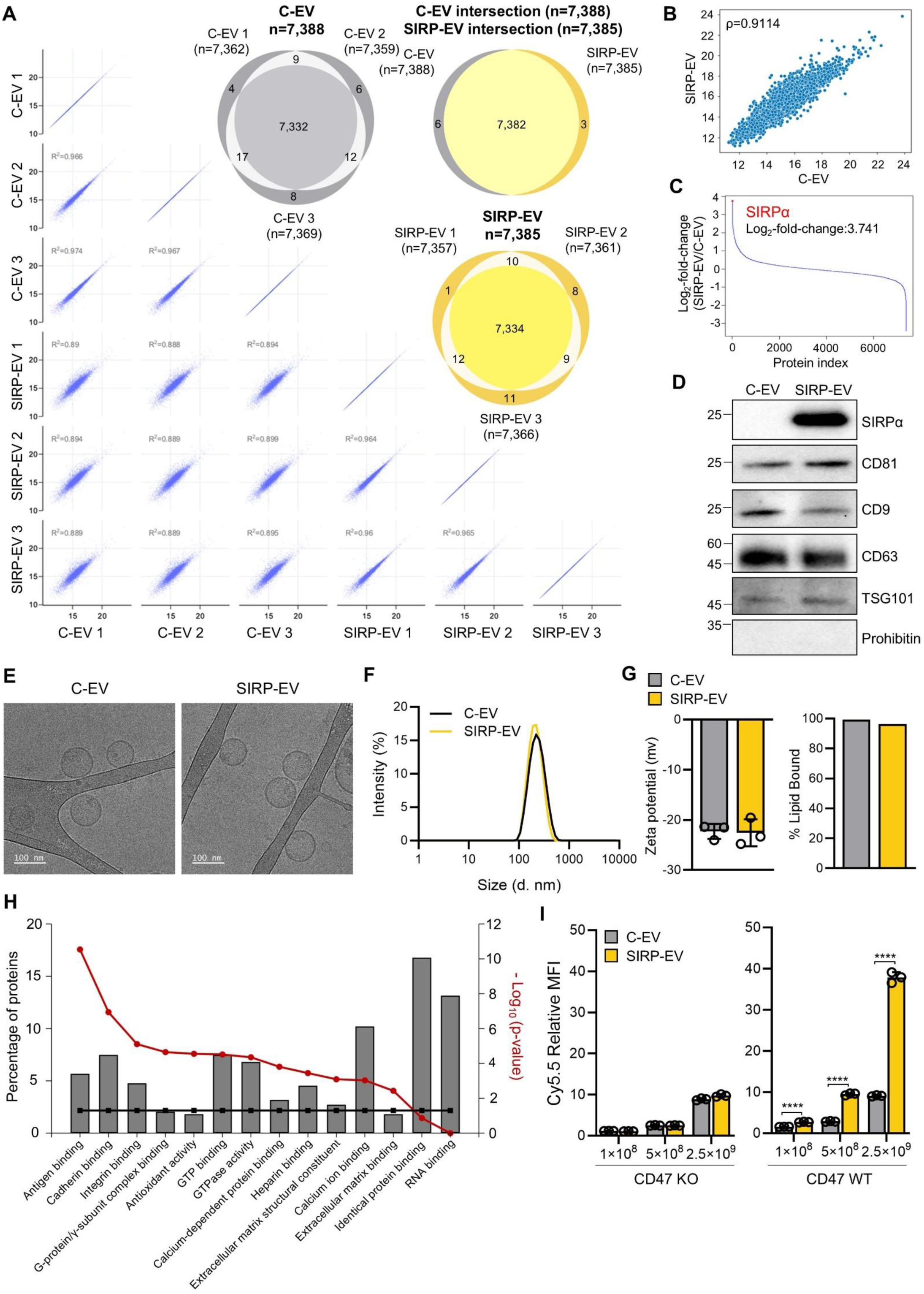
Comprehensive analysis of MSC-derived SIRP-EV. (A-C) Protein profile comparisons and reproducibility between C-EVs and SIRP-EVs. (A) Scatter plot consistency and R² values for protein identification across replicates; Venn diagram overlaps of EV type-specific proteins. (B) Correlation coefficient analysis of protein profile similarities between replicates. (C) Quantification of SIRPα overexpression in SIRP- EVs compared to C-EVs. (D) Protein expression of EVs to examine SIRPα, EV markers (CD81, CD9, CD63, and TSG101), and non-EV marker, prohibitin. (E) Representative cryo-TEM image of EVs. (F) Size distribution of EVs analyzed by DLS. (G) Zeta potential measurements of EVs analyzed using NTA, which denote the surface charge distribution (left, n=3). MemGlow staining of EVs (right). (H) Network diagram displaying over-represented Gene Ontology Biological Processes (GOBPs) among the top 506 SIRP-EV proteins. Analysis shows the proportion of these proteins compared to the total, with significant GOBPs (p < 0.05) indicated by black and red lines for group and individual function significance, respectively. (I) Quantification of cell binding, following a 30-min co-incubation of Cy5.5-labeled C-EV and SIRP-EV with CD47 KO and CD47 WT cells (n=3). Bar graph data are presented as mean ± S.D. Statistical significance was determined by two-tailed unpaired student’s t-test (G), two-way ANOVA with Sidak’s post-hoc test by comparing the groups treated with the same concentration of EVs (I). P****<0.0001.

To assess the targetability of SIRP-EVs toward CD47, we conducted *in vitro* binding assays using CD47 knockout (CD47 KO) and wild-type (WT CD47) cell lines. These assays showed that SIRP-EVs bind more selectively to WT CD47 cell line in a dose-dependent manner, compared to C-EVs, while no significant binding was observed in CD47 KO cell line **(Fig. 3I)**. Additionally, considering the known issues associated with CD47 antibodies—such as red blood cell (RBC) aggregation due to their Fc regions^21^—we carried out comparative *in vitro* hemagglutination assays between human CD47 antibodies and SIRP-EVs. Significantly, CD47 antibodies caused increased haziness indicating human RBC aggregation, while SIRP-EVs did not induce such adverse effects **(Supplementary Fig. 11A-C)**. Further toxicity evaluations of SIRP-EVs, through intravenous administration (i.v.) in mice at doses exceeding therapeutic levels, showed no observable toxicity **(Supplementary Fig. 11D)**. These results highlight SIRP-EVs as a promising alternative to traditional CD47 antibodies.

### Intravenous delivery of SIRP-EV demonstrates preferential accumulation in CD47- overexpressing injured liver tissue

We investigated the biodistribution of SIRP-EV, focusing on their accumulation based on CD47 expression. Cy5.5 dye-labeled SIRP-EVs were administered intravenously to both normal mice and models with APAP-ALF. Initial verification using fast protein liquid chromatography confirmed that Cy5.5 dye was solely associated with the SIRP-EVs, with no free dye detected **(Fig. 4A)**. In the APAP-ALF models, which overexpress CD47, these Cy5.5- labeled SIRP-EV accumulated more in the liver tissues of compared to those in normal mice **(Fig. 4B, C)**.

**Fig. 4:**
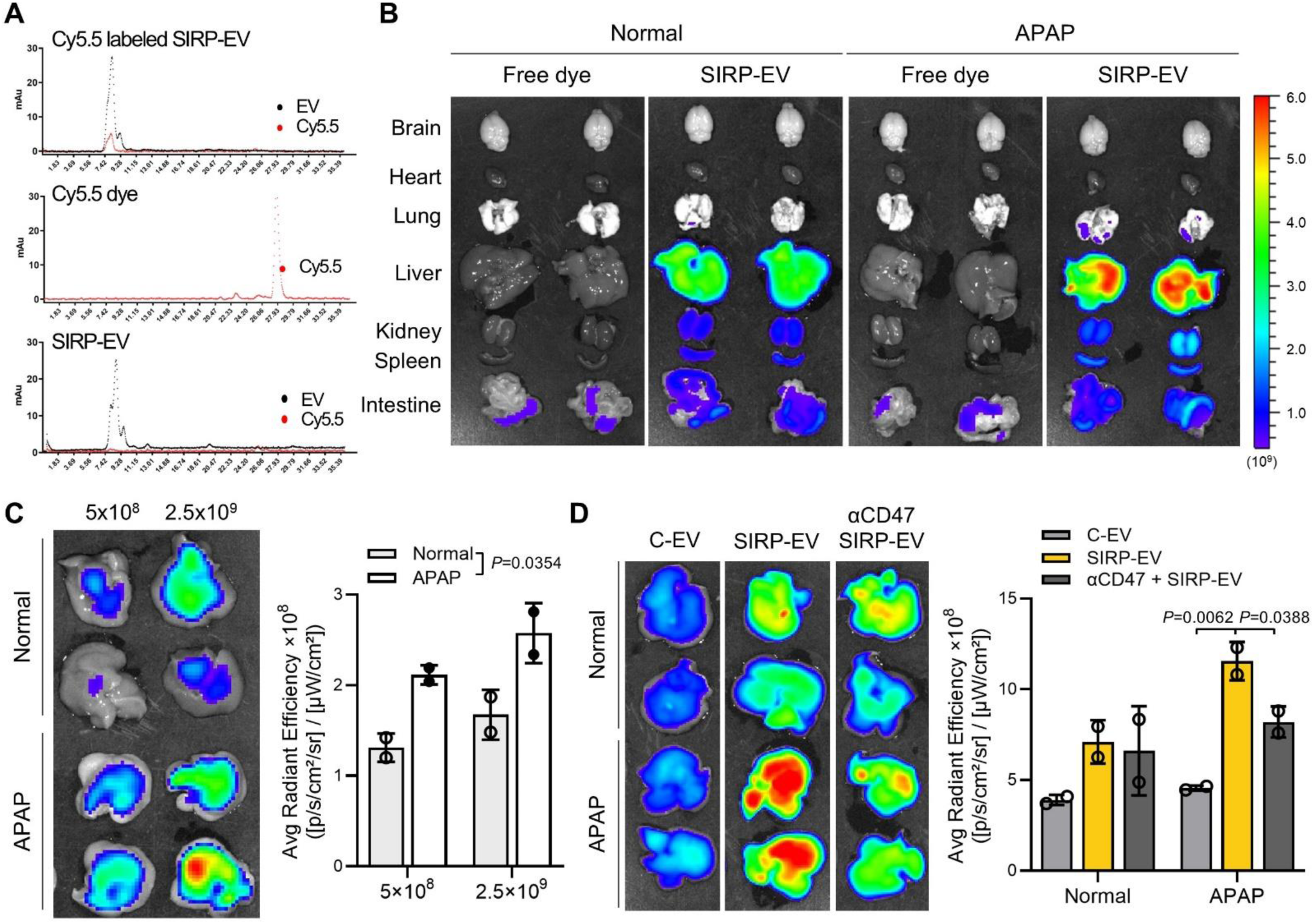
SIRP-EV preferentially accumulates in CD47-overexpressing injured liver after systemic delivery. (A) FPLC elution profiles of Cy5.5-labeled SIRP-EV, Cy5.5 dye, and SIRP-EV. Fluorescence was detected using a UV-Vis detector at 280 nm and 700 nm. (B) *Ex vivo* imaging of the organ distribution in mice 24 h after intravenous administration of 2.5 × 10^10^ Cy5.5-labeled SIRP-EVs. (C-D) Systemic delivery and uptake of EVs in mice liver tissues. (C) Dose-dependent accumulation of SIRP-EVs in liver tissues of APAP-ALF mice was observed within 24 h. Representative images of the *in vivo* imaging system (IVIS) are shown (left), and quantitative fluorescence signals from *ex vivo* livers are presented (right) (n=2). (D) A reduction in SIRP-EV accumulation in APAP-ALF liver tissues resulted from pre-blocking with the anti-CD47 antibody (n=2). Bar graph data are presented as mean ± S.D. Statistical significance was determined by two-tailed unpaired student’s t-test (C), and one-way ANOVA with Tukey’s post-hoc test (D).

We further confirmed the CD47-dependent biodistribution of SIRP-EV by pre-treating with CD47 antibodies, which resulted in decreased accumulation of SIRP-EV in CD47- overexpressing liver tissues **(Fig. 4D and Supplementary Fig. 12A)**. Intriguingly, SIRP-EVs displayed higher *in vivo* signals not only in the APAP-ALF model but also in normal mice 24 hours post-injection, compared to C-EVs. **(Supplementary Fig. 12A)**. Furthermore, the use of an isotype control antibody (IgG) showed no reduction in liver accumulation of SIRP-EVs, further supporting the CD47-dependent nature of SIRP-EV accumulation in injured liver tissue (**Supplementary Fig. 12B**). These findings collectively underscore the selective and CD47- dependent accumulation of SIRP-EVs in APAP-induced liver injury.

### SIRP-EV accumulates in CD47-overexpressing necroptotic hepatocytes and exhibits therapeutic efficacy in ALF

We assessed whether SIRP-EVs could accumulate in CD47-overexpressing necroptotic hepatocytes in ALF-induced liver tissue, promoting efferocytosis and liver regeneration. To do this, we conducted evaluations across various experimental schedules (**Supplementary Fig. 13**). Our results showed that SIRP-EV treatment significantly decreased CD47 expression in injured liver tissues (**Fig. 5A**). Furthermore, multiplex immunohistochemistry (IHC) provided deeper insights into CD47 expression dynamics within necrotic liver areas induced by APAP, which were primarily surrounded by macrophages (F4/80^+^) **(Fig. 5B)**. In contrast, liver tissues treated with SIRP-EVs exhibited dispersed macrophages, reduced necrotic areas, and lower CD47 expression, accompanied by a significant decrease in the ratio of CD47^+^ necroptotic cells **(Fig. 5C, D and Supplementary Fig. 14)**.

**Fig. 5:**
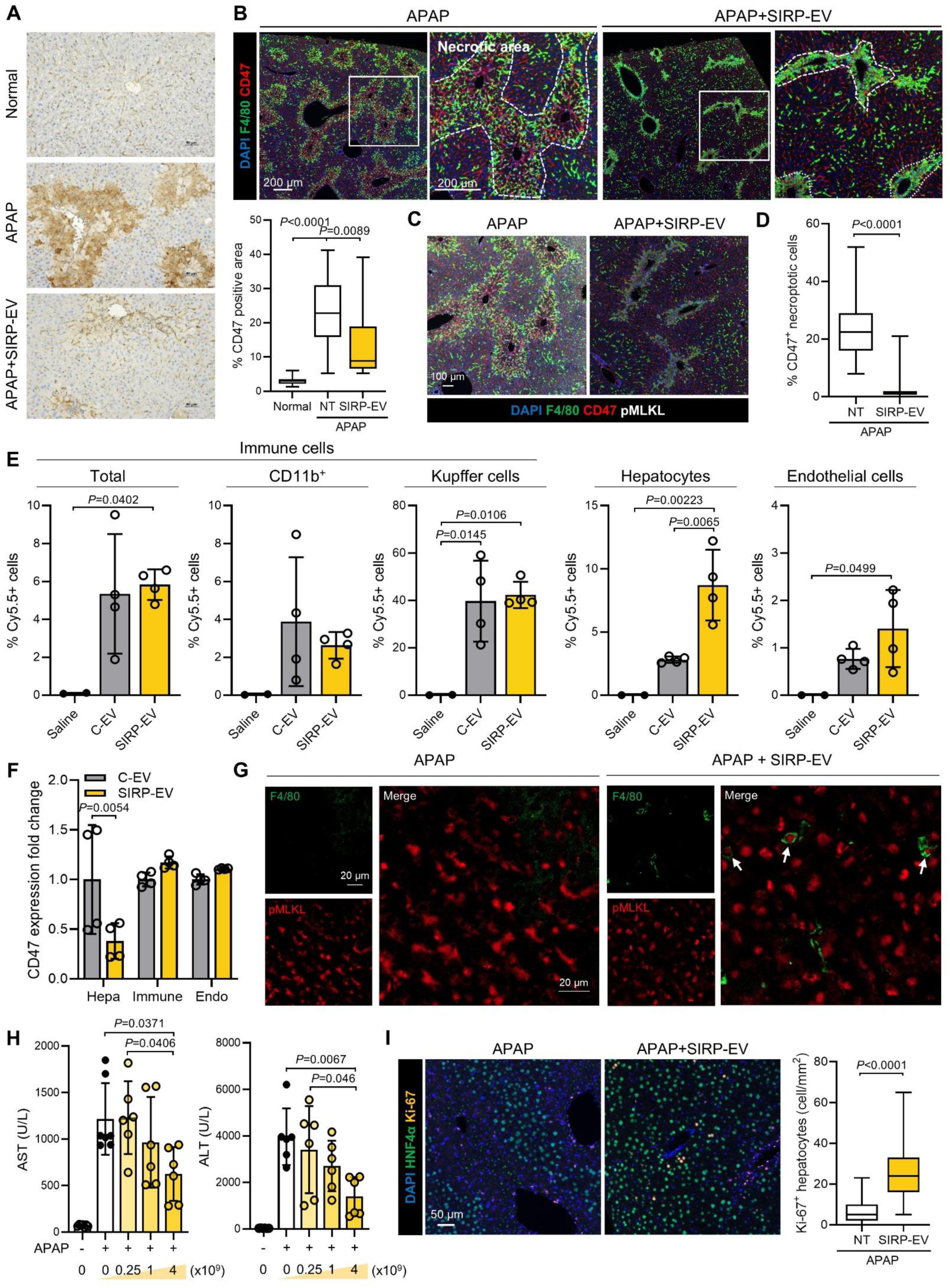
SIRP-EV targets CD47-expressing necroptotic hepatocytes to promote therapeutic efficacy via enhanced efferocytosis and liver regeneration in ALF. (A) Representative histological images of CD47 staining (left) and quantification of CD47-positive area (right) in liver tissue from normal and APAP-ALF mice (B) Representative multiplex IHC images from APAP-ALF mouse liver after 4 × 10^9^ SIRP-EVs treatment. The left panel shows a broader view, while the right panel focuses on a necrotic area, as delineated by the dashed line. (C) Representative multiplex IHC of APAP-ALF mouse liver tissue and (D) the quantification of CD47-positive necroptotic cells (CD47^+^pMLKL^+^) is shown on the right panel. (E) Cell type-specific biodistribution of SIRP-EV in APAP-ALF livers (n=2 or 4 per group). (F) CD47 expression in liver tissue cell types after systemic treatment with SIRP-EVs compared to C-EVs (n=4). (G) Representative confocal images of liver sections from mice 24 h post-induction with a 500 mg/kg APAP dose, treated with 4 × 10^9^ SIRP-EVs. Cells were immunostained for F4/80 (green) and pMLKL (red), with white arrowheads indicating engulfed necroptotic cells. (H) Serum AST and ALT levels were determined 48 h after APAP injection in subjects receiving the indicated dose of SIRP-EV (n=6 or 7 per group). (I) Representative multiplex IHC of APAP-ALF mouse liver treated with 4 × 10^9^ SIRP-EVs. Quantification of Ki-67 positive proliferative hepatocytes (Ki67^+^HNFα^+^) in APAP-ALF mouse liver is shown in the right panel. Bar graph data are presented as mean ± S.D. Statistical significance was determined by one-way ANOVA with Tukey’s post-hoc test (A, E, and H), two-tailed unpaired student’s t-test (D and I) and two-way ANOVA with Sidak’s post-hoc test (F). Hepa, Hepatocytes; Immune, Immune cells; Endo, Endothelial cells.

To confirm SIRP-EV accumulation in necroptotic hepatocytes, we administered fluorescently labeled EVs systemically in the APAP-ALF model and used flow cytometry to evaluate cellular uptake across cell types (**Supplementary Fig. 15A**). Myeloid cells, particularly Kupffer cells, are critical immune cells for the primary defense within liver tissue^10,22^. Kupffer cells are generally known to predominantly accumulate EVs within the liver^23^. Consequently, we identified that over 40% of Kupffer cells had EVs distributed within them. This result was consistent regardless of SIRPα expression on the EVs **(Fig. 5E)**. Notably, SIRP-EVs appeared to reach hepatocytes more effectively compared to C-EVs **(Fig. 5E)**. Furthermore, we observed that the accumulation of SIRP-EVs in hepatocytes was abolished when the cells were pre-treated with CD47-blocking antibodies (**Supplementary Fig. 15B**). These findings indicate that systemically administered SIRP-EVs achieve enhanced targeting of necroptotic hepatocytes that overexpress CD47, surpassing the targeting capabilities of C-EVs.

Interestingly, CD47, which was overexpressed in hepatocytes of the APAP-ALF model, showed a significant decrease following SIRP-EV treatment (**Fig. 5F and Supplementary Fig. 16**). To determine whether this CD47 reduction facilitated macrophage efferocytosis of necroptotic hepatocytes, we performed tissue staining and analyzed liver sections via confocal microscopy. Our findings revealed that SIRP-EV treatment significantly increased the presence of macrophages (F4/80^+^) and enhanced efferocytosis of necroptotic cells (pMLKL^+^) in liver tissues of APAP-ALF model. **(Fig. 5G)**. Supporting this result, bone marrow-derived macrophages (BMDMs) exhibited enhanced phagocytosis of APAP-treated HepG2 cells and primary hepatocytes from APAP-ALF models incubated with SIRP-EVs compared to C-EVs (**Supplementary Fig. 17**).

Finally, we evaluated the therapeutic efficacy of SIRP-EVs in the APAP-ALF model. SIRP-EV administration led to a dose-dependent reduction in liver toxicity markers, such as AST and ALT (**Fig. 5H**). Additionally, a single injection of SIRP-EV in the APAP-ALF model resulted in an increase in Ki67^+^ hepatocytes, indicative of liver regeneration (**Fig. 5I**). These findings suggest that SIRP-EVs not only bind to and suppress CD47 on necroptotic hepatocytes but also promote macrophage efferocytosis, resulting in enhanced therapeutic efficacy in a dose-dependent manner.

### Single injection of SIRP-EV demonstrates therapeutic efficacy across multiple ALF models

To investigate the enhanced therapeutic efficacy mediated through SIRPα in our engineered EVs compared to C-EVs, we assessed their therapeutic potential in various ALF models (**Supplementary Fig. 13**). In the APAP-ALF model, histological analysis revealed that SIRP- EV treatment significantly reduced dead cell counts in damaged liver tissues (**Fig. 6A, B**). Treatment with SIRP-EVs also resulted in a higher ratio of monocyte-derived macrophages (MoMF) to the Ly6C low type and Kupffer cells, crucial for inflammation resolution, compared to C-EVs **(Fig. 6C and Supplementary Fig. 18A-C)**. Additionally, SIRP-EVs were associated with reductions in pro-inflammatory cytokine IL-6 levels and neutrophil infiltration, which are indicative of acute inflammation **(Supplementary Fig. 18D, E)**. Moreover, SIRP-EV treatment led to statistically significant improvements in liver toxicity markers, AST and ALT, surpassing C-EVs in the APAP-ALF model **(Fig. 6D)**.

**Fig. 6.**
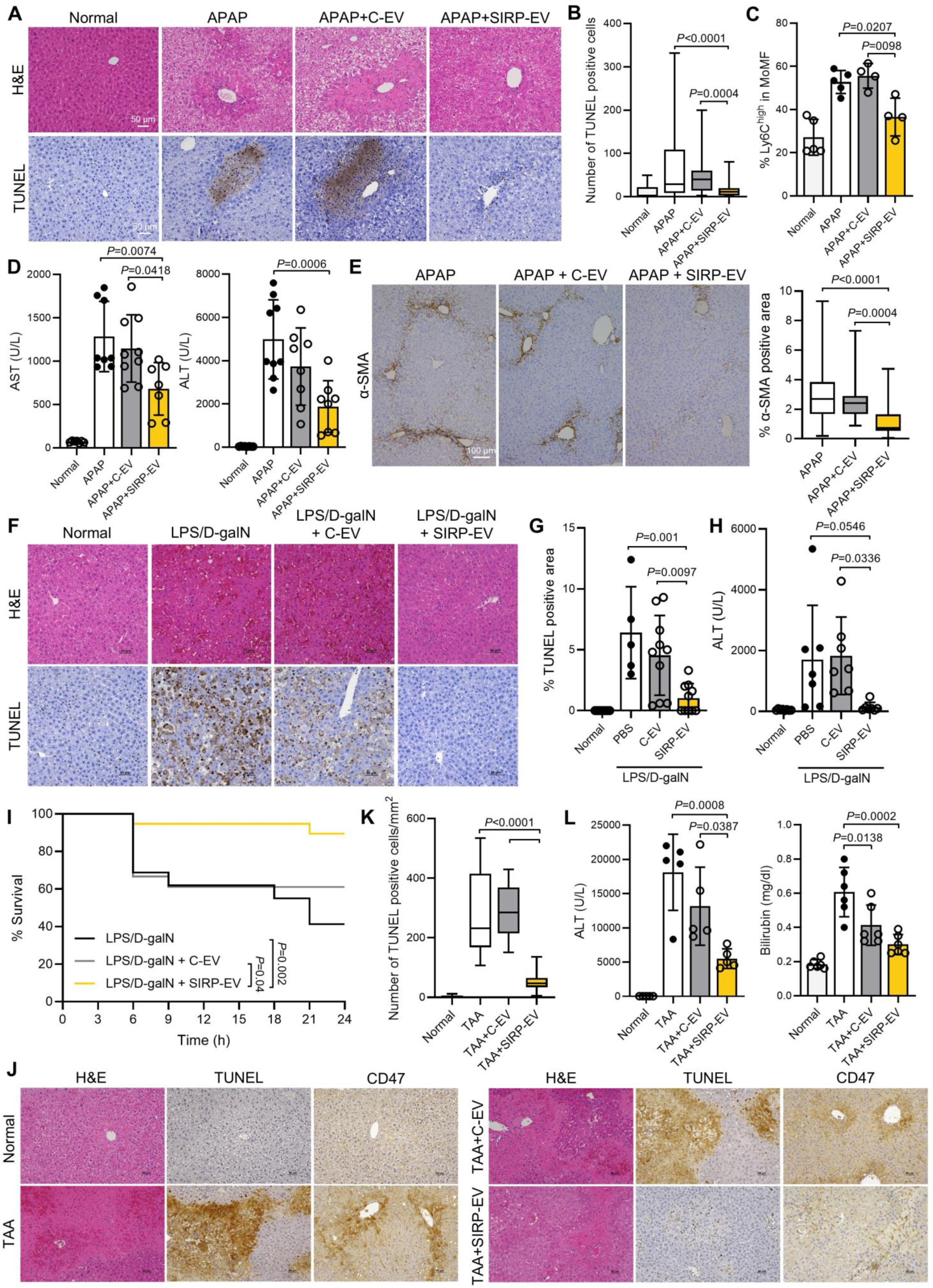
SIRP-EV outperforms C-EV in ALF models, attributed to SIRPα expression. (A) Representative histological images of H&E and TUNEL in liver tissue from normal and APAP-ALF mice after EVs treatment. (B) Quantification of TUNEL-positive cells. (C) Flow cytometry determination of Ly6C^high^ as a percentage of total liver CD11b^high^ F4/80^low^ MoMFs (n=4 or 5 per group). (D) Serum AST and ALT levels were measured 48 h post-500 mg/kg APAP induction in subjects treated with 4 × 10^9^ EVs (n=7 to 9 per group). (E) Representative IHC images of α-SMA staining in liver tissues from mice that survived 120 h after induction with a 700 mg/kg dose of APAP. Quantification of α-SMA is shown in the right panel. (F-I) Therapeutic effects of 9 × 10^8^ EVs on an LPS/D-galN-induced ALF model. (F) Representative histological images of H&E and TUNEL staining in liver tissue from normal and LPS/D-galN-induced ALF mice after the indicated treatment. (G) Quantification of the TUNEL-positive area. (H) Serum ALT levels were determined 8 h post-LPS/D-galN injection (n=7 or 8 per group). (I) Kaplan–Meier survival curves for LPS/D-galN-induced ALF mice treated with EVs (n=16 to 19 per group, across three independent experiments). (J-L) Therapeutic efficacy of 5 × 10^8^ EVs in TAA-induced ALF models. (J) Representative histological images of H&E, TUNEL, and CD47 staining in liver tissue from both normal and TAA-induced ALF mice following EVs treatment. (K) Quantification of TUNEL-positive cells in liver tissue. (L) Serum ALT and total bilirubin levels were determined 40 h post-TAA injection (n=5 to 6 per group). Bar graph data are presented as mean ± S.D. Statistical significance was determined by one-way ANOVA with Tukey’s post-hoc test (B-E, G, H, K, and L).

Notably, SIRP-EV treatment resulted in higher survival rates in the high dose APAP (700 mg/kg)-ALF model compared to C-EV and CD47 antibody treatments **(Supplementary Fig. 19A, B)**. Given that ALF progression is often linked to liver fibrosis^24^ and non-efferocytosed damaged hepatocytes can induce fibrogenic activation^25^, we investigated whether SIRP-EVs could prevent fibrosis by facilitating the clearance of these damaged hepatocytes. Our findings revealed that SIRP-EV treatment notably mitigated liver fibrosis compared to C-EVs **(Fig. 6E)**, underscoring the effectiveness of SIRP-EVs in targeting CD47^+^ necroptotic hepatocytes and broadening their potential applications in preventing ALF complications, including fibrosis.

In the LPS/DgalN-induced ALF prevention model, SIRP-EVs not only improved survival rates but also substantially reduced ALT levels and the ratio of dead cells in liver tissues more effectively than C-EVs **(Fig. 6F-I and Supplementary Fig. 19C)**. Similarly, in the TAA- induced ALF model, SIRP-EVs exhibited superior therapeutic effects compared to C-EVs, as evidenced by lower dead cell ratios, reduced CD47 expression, and decreased serum ALT and bilirubin levels **(Fig. 6J-L and Supplementary Fig. 19D)**. Overall, these results indicate that incorporating SIRPα into C-EVs significantly enhances their therapeutic efficacy.

### SIRP-EV enhances therapeutic efficacy by transferring MSC protein cargos in ALF

We further verified that the therapeutic efficacy of SIRP-EV originates from their MSC- derived cargos by comparing them with SIRP-EVs isolated from HEK cells. We genetically modified HEK cells to express SIRPα, producing HEK cell-derived SIRP-EV(_HEK_SIRP-EV), and evaluated their efficacy in ALF models. SIRP-EVs were more effective than _HEK_SIRP-EVs, as demonstrated by a significant reduction in blood levels of AST and ALT **(Fig. 7A)**. Furthermore, histological analysis demonstrated that SIRP-EVs significantly decreased dead cell counts and neutrophil infiltration compared to _HEK_SIRP-EV **(Fig. 7B, C)**. This suggests that the enhanced therapeutic effects of SIRP-EV are not solely due to CD47 blocking but also involve other contributory factors.

**Fig. 7.**
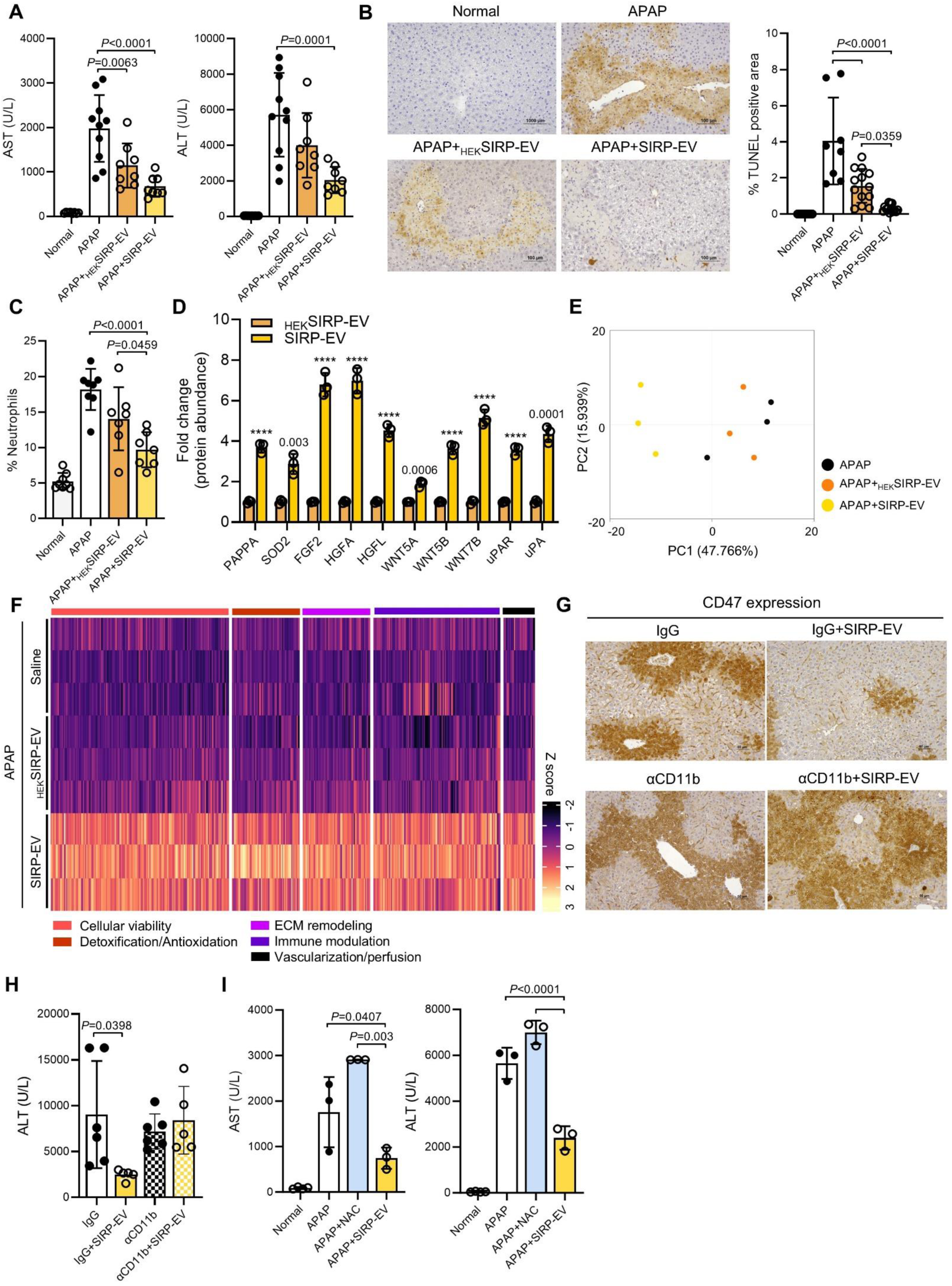
SIRP-EV leverages MSC properties to elevate ALF treatment, promoting regeneration and diminishing inflammation. (A-F) Therapeutic efficacy of 4 × 10^9^ EVs in 500mg/kg APAP-ALF models. (A) Serum AST and ALT levels were determined 48 h after APAP injection in subjects treated with EVs (n=8 to 10 per group). (B) Representative histological images of TUNEL staining in liver tissue from normal and APAP-ALF models after the indicated treatment. Quantification of the TUNEL-positive area is shown in the right panel. (C) Flow cytometry determination of Ly6G-positive neutrophils as a percentage of the total liver CD45^+^ leukocytes (n=7 or 8 per group). (D) The abundance of proteins associated with the prevention of cell death (PAPPA, SOD2), cell proliferation (FGF, HGF, WNT ligands), and angiogenesis (uPAR, uPA) in SIRP-EVs compared to _HEK_SIRP- EV (n=3 per group). P-values are annotated on the bar, ****P < 0.0001. (E) Principal component analysis (PCA) of RNA sequencing data from CD11b^+^ cells from liver tissues in APAP, APAP+ _HEK_SIRP-EV, and APAP+SIRP- EV groups (n=3 per group). (F) Heatmap of 404 representative genes from each gene cluster identified. (G and H) Therapeutic efficacy of 4 × 10^9^ SIRP-EVs in APAP-ALF models with CD11b reduction. (G) Representative histological images showing CD47 staining in liver tissue from APAP-ALF models following the specified treatment. (H) Serum ALT levels were measured 48 hours post-APAP injection in subjects receiving the indicated treatments (n=5 to 6 per group). (I) Serum AST and ALT levels were measured 48 h after injection of 300 mg/kg APAP-ALF models in subjects receiving the indicated treatments (n=3 or 4 per group). Bar graph data are presented as mean ± S.D. Statistical significance was determined by one-way ANOVA with Tukey’s post-hoc test (A-C, H and I), two-tailed unpaired Student’s t-test (D), Benjamini-Hochberg adjusted P-values (F).

We next analyzed and compared the functional protein cargos known to promote regeneration^20^ in both types of EV, aiming to elucidate the differences in therapeutic efficacy between MSC-derived SIRP-EVs (SIRP-EVs) and HEK cell-derived SIRP-EVs (_HEK_SIRP- EVs). The analysis revealed that proteins associated with preventing cell death (PAPPA, SOD2), promoting cell proliferation (FGF, HGF, WNT ligands), and enhancing angiogenesis (uPAR, uPA) were more abundantly present in SIRP-EVs than in _HEK_SIRP-EVs **(Fig. 7D)**.

Acute and severe liver injury drives substantial migration of myeloid cells into the liver, where they differentiate into macrophages that play essential roles in both exacerbating liver injury and promoting tissue repair alongside resident macrophages, such as Kupffer cells^19,26^. Given our biodistribution data demonstrating that SIRP-EVs were effectively distributed within myeloid cells (CD11b^+^ cells) and Kupffer cells (CD11b^+^F4/80^high^ cells), we analyzed the regenerative efficacy of SIRP-EVs in these cells. Notably, transcriptomic analysis of liver CD11b^+^ cells in liver tissues from ALF models indicated that a single dose of SIRP-EV systemic treatment showed distinct genetic differences compared to both the _HEK_SIRP-EV and non-treated groups **(Fig. 7E)**. Heat map analysis of 404 statistically differentially expressed genes among the three groups demonstrated that SIRP-EV treatment significantly upregulated genes associated with cellular viability, detoxification/antioxidation, extracellular matrix (ECM) remodeling, immune modulation, and vascularization/perfusion, all of which are related to tissue regeneration **(Fig. 7F)**.

To further elucidate whether the observed therapeutic effects of SIRP-EVs in ALF models are dependent on liver CD11b^+^ cells, we performed experiments using CD11b neutralizing antibodies (**Supplementary Fig. 13**). Consistent with previous studies^26^, ALF models showed a marked increase in CD11b^+^ cells in the liver and spleen compared to normal conditions (**Supplementary Fig. 20**). Treatment with CD11b neutralizing antibodies effectively reduced CD11b^+^ cell levels in the spleen to near-normal and significantly decreased liver CD11b^+^ cells relative to untreated APAP-ALF models, although levels remained higher than normal livers (**Supplementary Fig. 20**). Intriguingly, APAP-ALF models treated with the CD11b neutralizing antibody did not exhibit the reduction in serum ALT levels or the decreased hepatic CD47 expression observed in SIRP-EV-treated mice that received control IgG (**Fig. 7G, H**). These findings suggest that the therapeutic efficacy of SIRP-EVs in ALF may rely on macrophages, potentially not only by removing necroptotic hepatocytes through CD47 blockade but also by enhancing the regenerative capacity in liver tissue via MSC-derived cargo.

Currently, N-acetylcysteine (NAC) is the standard treatment for ALF caused by APAP overdosing in clinical settings, primarily preventing hepatocyte death^27^. However, its effectiveness depends on the timing of APAP exposure. Previous studies have shown that NAC loses its protective effects if administered more than 3 hours after APAP exposure^28^. A comparative efficacy study between SIRP-EV and NAC was conducted by administering both therapeutics 8 hours after APAP injection. The results demonstrated that SIRP-EV offered superior therapeutic benefits **(Fig. 7I and Supplementary Fig. 21)**. This finding underscores the potential of SIRP-EV as a competitively advantageous alternative to current ALF treatments.

## Discussion

Acute inflammation impairs the resolution capabilities of immune cells and diminishes their inherent regenerative capacities, necessitating sophisticated therapeutic approaches^29^. This study introduces a dual-mode action therapeutic strategy utilizing SIRP-EVs derived from engineered MSCs. These SIRP-EVs not only enhance the phagocytic activity of liver macrophages to clear necroptotic hepatocytes via CD47 blockade but also promote liver regeneration by delivering MSC-derived cargo that reprograms macrophages to support tissue repair **(Fig. 8)**.

**Fig. 8.**
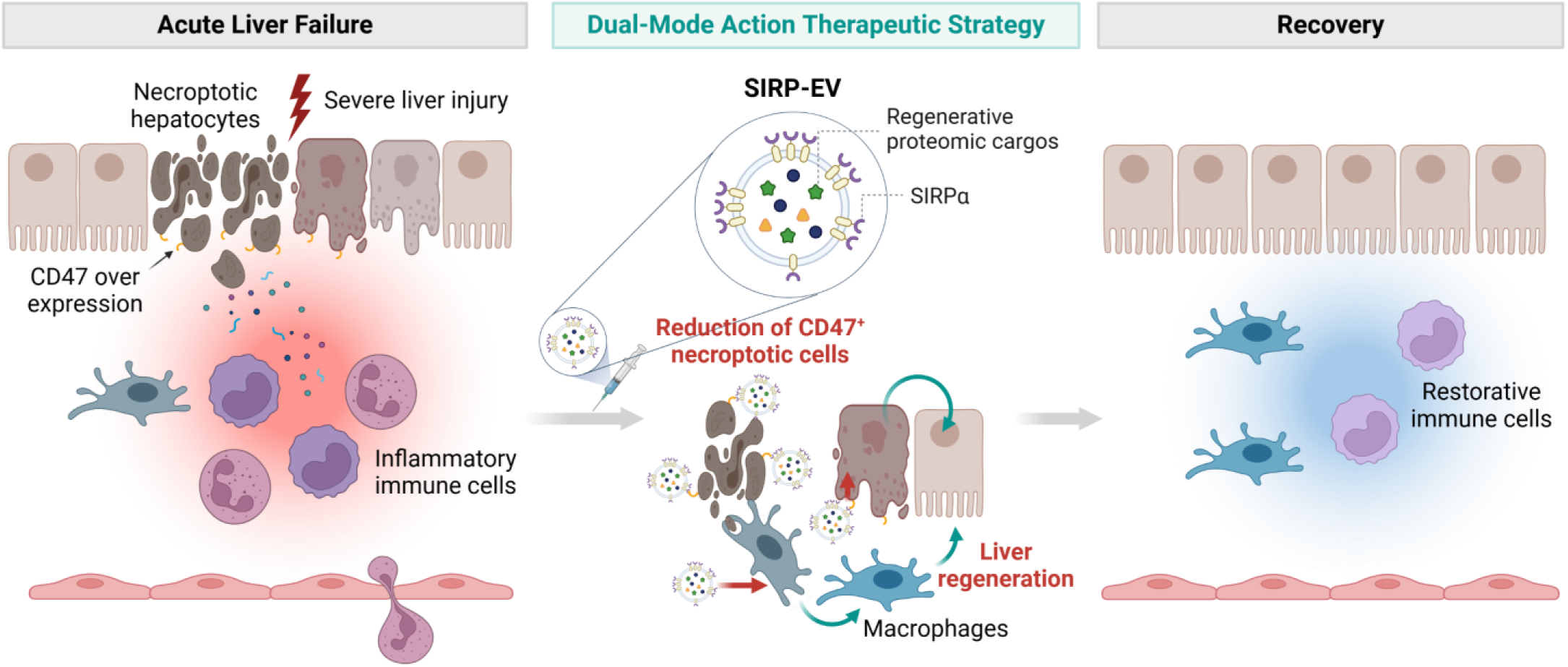
A dual-mode action therapeutic strategy for ALF: SIRP-EV from engineered mesenchymal stem cells resolves CD47 in necroptotic hepatocytes and delivers regenerative cargo. SIRP-EVs mitigate necroptosis by targeting CD47 on necroptotic hepatocytes, modulate the function of macrophages, and enhance hepatocyte regeneration, ultimately promoting liver regeneration in ALF.

As embryonically derived tissue-resident macrophages, Kupffer cells occupy niches that limit the differentiation of circulating CD11b^+^ monocytes into macrophages under homeostatic conditions^30^. However, under acute inflammation or severe injury, resident Kupffer cells become largely depleted, allowing CD11b^+^ monocytes from the bone marrow to migrate and repopulate the vacant niches^30^. These MoMF precursors possess an almost identical potential to develop into Kupffer cells and have been reported to play a critical role in both exacerbating and resolving inflammation during hepatic injury^30,31^. Unlike embryonic Kupffer cells, CD11b^+^ monocyte-derived Kupffer cells exhibit greater resistance to apoptosis and possess enhanced phagocytic and tissue repair capacities^32^.

In our ALF model, we observed a substantial depletion of Kupffer cells. However, SIRP-EV treatment increased the overall proportion of Kupffer cells and induced a shift of MoMFs toward a resolutive Ly6C low phenotype. Furthermore, SIRP-EVs promoted the clearance of CD47-overexpressing necroptotic hepatocytes by liver macrophages and significantly upregulated genes associated with tissue regeneration in liver CD11b^+^ cells. The observed reduction in therapeutic efficacy in a CD11b-diminished ALF model supports the hypothesis that CD11b^+^ cells are integral to SIRP-EV’s therapeutic mechanism. Our study does not definitively clarify whether the primary effectors in necroptotic cell clearance are embryonically derived or CD11b^+^ monocyte-derived macrophages, nor does it specify the roles of different CD11b^+^ cell types, such as monocytes, dendritic cells, and granulocytes. Future studies are needed to elucidate the specific cellular contributors to SIRP-EV’s therapeutic effects in liver regeneration and inflammation resolution.

Recent studies have identified necroptotic hepatocytes as a significant contributor to the exacerbation of ALF^6^. This study is the first to demonstrate that these necroptotic hepatocytes overexpress CD47 in ALF. Our findings indicate that blocking CD47 with SIRP-EVs reduces necroptotic hepatocytes and normalizes CD47 expression, suggesting its utility as a biomarker for ALF and potentially other diseases. Further support for this hypothesis is provided by spatial transcriptomic analysis, which reveals significant increases in SIRPα expression at sites of necroptosis and macrophage interaction. These results reinforce the potential of CD47 and SIRPα as biomarkers in severe acute inflammatory diseases.

The framework for EV production and purification has been established in previous studies^33–36^. This study demonstrates a robust method for producing high-purity, high-yield SIRP-EVs while preserving the intrinsic properties of MSCs. To accelerate the clinical translation of SIRP-EVs, we optimized a scalable manufacturing process to maximize EV yield using a 3D bioreactor system. Additionally, we developed a downstream process to concentrate EVs and remove non-EV impurities, such as nucleic acids released from cells and media proteins (e.g., albumin), to meet regulatory expectations^37^. This process achieved 93 % and 99 % reductions in nucleic acids and albumin, respectively, through filtration and chromatography, while maintaining high EV recovery. Using the particle-to-protein ratio (p/mg protein) as a purity metric^38,39^, we improved the ratio from 7.5E+10 to 8.2E+11 (∼11-fold increase) and achieved a 50% recovery rate. This efficient, cost-effective approach offers a promising platform for developing novel therapeutics using engineered EVs derived from MSCs.

Interestingly, the EVs produced by this method contained significantly low RNA content compared to those isolated through traditional research-grade EV production processes. While RNA is generally thought to be encapsulated within EVs, our findings suggest that RNA may be associated with the outside of EVs, warranting further investigation. Although numerous studies confirm the biological effects of miRNAs in MSC derived EVs^17^, the RNA amount of approximately 1 ng in the SIRP-EVs used for our *in vivo* experiments challenges the notion that the therapeutic efficacy we observed can be explained by these RNAs. Therefore, our study shifts focus to the regenerative potential of MSC protein cargos but emphasizes that additional functional analyses of other MSC-derived cargos, such as DNA, RNA, and lipids are necessary.

EVs are increasingly recognized as promising drug delivery systems, noted for their high biocompatibility and ability to cross biological barriers^40–43^. However, they primarily accumulate in liver macrophages^23^. To address this issue, recent studies have enhanced EV targeting by forming a protein corona on their surface, specifically coating C-EVs with albumin to improve delivery to hepatocytes beyond macrophages^23^. Additionally, surface engineering of EVs has been used to enhance targeted delivery to specific cells, thereby enhancing therapeutic efficacy^44–47^. Our findings prove that SIRPα expression on EVs not only mitigates necroptosis but also promotes effective distribution in CD47-overexpressing liver tissues and hepatocytes extending beyond macrophages in acute inflammatory conditions. This unique distribution of SIRP-EVs facilitates the delivery of MSC-derived regenerative factors and aids in blocking CD47 overexpression, which is observed in necroptotic cells. CD47 is not only overexpressed on pathological cells but also present on normal cells, posing challenges for therapies using CD47 antibodies^48,49^. These antibodies can inadvertently bind to normal cells, leading to unwanted clearance by immune cells that recognize the Fc region of antibodies. This limitation has narrowed the therapeutic window and contributed to clinical failures^50,51^. Importantly, SIRP-EVs, which lack Fc regions, do not present adverse side effects related to normal cell clearance.

Our research further indicates that even in normal mice, SIRP-EVs exhibit a prolonged *in vivo* presence compared to C-EVs. This finding implies that the presence of SIRPα on the EV surface may enhance their in *vivo* retention through interactions with CD47, which is expressed under normal conditions. Additionally, previous studies have shown that SIRP-EVs bind to RBCs via CD47, and these RBC-bound SIRP-EVs can subsequently transfer to cancer cells that overexpress CD47^21^. This implies that RBCs may act as carriers, promoting circulation and enhancing targeted delivery to pathological sites. While these findings provide a strong basis, further validation through additional replicates of the biodistribution experiments will be important. More detailed and comprehensive pharmacokinetic studies will be necessary to demonstrate the prolonged circulation and targeted delivery of SIRP-EVs.

Studies of liver fibrosis in ALF have been limited due to the severity of clinical syndrome^52,53^. However, some research has demonstrated that delayed clearance of dead hepatocytes and subsequent regeneration after injury can amplify the fibrotic response^25^. We have shown that SIRP-EV not only has therapeutic efficacy in ALF but also provides evidence for a preventing effect on fibrosis progression after acute injury. While promising anti-fibrotic effects have been observed in experimental animal models, further mechanistic and fibrosis-related investigations are necessary to replicate these results in clinical trials.

In conclusion, this study introduces a dual-mode action approach utilizing SIRP-EVs for both inflammation resolution and regeneration promotion. Additional efforts are underway to improve our understanding, and further investigations are needed to definitively demonstrate how different cell types respond to SIRP-EV-mediated EV-cargo release. Although the primary focus has been on ALF, this strategy holds potential for application across a range of inflammatory conditions. Additionally, the scalable and high-purity production process of these engineered MSC-derived EVs establishes a solid foundation for further research and development. This groundwork aims to facilitate clinical translation, potentially revolutionizing the management of acute inflammatory diseases.

## Methods

### ALF animal models

Male C57BL/6N mice (7-week-old) were purchased from Orient Bio (South Korea). The mice received intraperitoneal (i.p.) injections of APAP (300mg/kg, 500 mg/kg or 700 mg/kg, Sigma-Aldrich, A7085) after 16 hours of fasting and then were returned to a normal diet. For LPS/D- galN, TAA, and Carbone tetrachloride (CCl4) inducing ALF models, male C57BL/6N mice (7- old-week) received i.p. injection of LPS (10 µg/kg, Sigma-Aldrich, L2630) and D-galN (700 mg/kg, Sigma-Aldrich, G0500) simultaneously, TAA (300 mg/kg, Sigma-Aldrich, 163678) and CCl4 (1 mL/kg, Sigma-Aldrich, 289116). To treat ALF models, the mice received intravenous (i.v.) injections of EVs (0.25 × 10^9^ ∼ 4 × 10^9^ EV particles) via the tail vein, 8 hours after APAP induction, 2 hours before LPS/D-galN induction, and 16 hours after TAA induction. N- acetylcysteine (300 mg/kg, Sigma-Aldrich, A7250, i.v.) and anti-CD47 antibodies (10 mg/kg, BioXCell, BE0007, i.p.) were administered 8 hours following APAP induction. All mice were maintained in the pathogen-free animal research facility of the Korea Institute of Science and Technology (KIST), and all *in vivo* experiments were performed under the guidelines of the Institutional Animal Care and Use Committee (IACUC, KIST-IACUC-2021-154-5).

### Biochemistry, cytokine, hematology, and coagulation analysis

For biochemistry analysis (AST, ALT, ALP, Albumin, Total-bilirubin), the whole blood was collected through retro-orbital bleeding with micro-Hematocrit capillary tubes into EDTA tubes (Gsmeditech, 123105) for plasma, and serum tubes (Gsmeditech, 111205) for serum. To remove cells and clots, the blood in collection tubes was centrifuged at 2,000 g for 20 min, once for serum and twice for plasma. The biochemistry was analyzed using AU480 Chemistry Analyzer (Beckman Coulter). For cytokines analysis, serum was collected as described above and IL-6 was measured using the mouse IL-6 Quantikine ELISA kit (R&D System, M600B- 1). For hematology and coagulation analysis, the whole blood was collected through cardiac puncture and blood samples were distributed into appropriate tubes [EDTA tube (BD microtainer, BD365974) for hematology (RBC, hematocrit, hemoglobin) analysis and trisodium citrate 0.109M (V-TUBETM, 630187) for coagulation analysis (PT, aPTT)] and analyzed by Sysmex XN-9000 (Sysmex) for hematology and CA-1500 (Sysmex) for coagulation test.

### Immunohistochemistry

The paraffin blocks of liver tissues fixed with 4% formalin were prepared, and sliced tissue samples were stained through Bond-X automatic slide stainer (Vision Biosystem) with primary antibody; CD47 (GeneTex GTX53912), RIP3 (Affinity Biosciences, AF7942), pMLKL (Affinity Biosciences, AF7420). The CD47 positive area and fibrosis (α-SMA) were quantified as a percentage of the stained area relative to the whole area of samples. TUNEL-stained samples were analyzed to quantify TUNEL-positive cell area relative to the total sample area.

### EV biodistribution

EVs were labeled with sulfo-Cyanine5.5 NHS ester or sulfo-Cyanine5.5 carboxylic acid (Lumiprobe, 27390) to assess the biodistribution of the EVs. Each EV was labeled with Cy5.5 dyes at a ratio of 5 x 10^10^ particles per 1 μg of Cy5.5 dye and incubated at 4°C overnight. Zeba™ Spin Desalting Columns (ThermoFisher, A57762) were utilized to purify the labeled EVs and eliminate free Cy5.5 dyes. The fluorescence intensity of Cy5.5-NHS conjugated EVs was analyzed using a microplate. The efficiency of Cy5.5 labeling was determined using Fast Protein Liquid Chromatography (FPLC) (Cytiva, AKTApurifierTM) and Superdex 200 Increase 10/300 GL columns (Cytiva, GE28-9909-44). To assess the biodistribution of EVs, 7- week-old male C57BL/6 mice were utilized for the APAP-ALF model analysis. 8 h post-APAP induction, Cy5.5-labeled samples at concentrations of 5 × 10^8^, 2.5 × 10^9^, or 2.5 × 10^10^ were administered intravenously. To assess the CD47-mediated targeting effect of SIRP-EVs, anti-CD47 antibodies (BioXCell, BE0007) and IgG2a isotype control antibodies (BioXCell, BE0090) (10 mg/kg, i.p.) were injected 30 min prior to the administration of 2.5 x 10^10^ Cy5.5- labeled EVs. After 24 hours in the injection of EVs, organs were excised for *ex vivo* analysis to determine organ accumulation. The *ex vivo* biodistribution of EVs was visualized using the IVIS Spectrum (Caliper Life Sciences, IVIS^®^ Lumina Series III). The diet of the mice was completely switched to alfalfa-free feed. For the evaluation of a single-cell-based EV distribution assay, liver tissues from the ALF model were dissociated using the gentleMACS™ Octo Dissociator with Heaters (Miltenyi Biotec, 130-096-427). Subsequently, individual cell populations, including hepatocytes, endothelial cells, myeloid cells, Kupffer cells, and MoMFs, were isolated in accordance with the liver tissue single-cell dissociation method. Further, the isolated single cells were labeled with an antibody panel of flow cytometry, and the Cy5.5 signals from each single cell were detected by CytoFLEX flow cytometry.

### Engineered MSC production

Human bone marrow-derived MSC (hBM-MSC) master cell banks (Passage 0, RoosterBio) were plated in RoosterNourish^TM^ (RoosterBio) in T-75 flasks (Corning) and incubated at 37°C, 5% CO_2_. After 2 days, cells were harvested and counted for seeding to the T-75 flasks. To engineer EVs displaying SIRPα, we generated a SIRPα variant capable of binding both human and mouse CD47 with high affinity^21,40,43,54^, fused to a type 1 transmembrane domain to ensure effective surface expression on EVs^55^. Lentiviral transduction was performed in RoosterGEM™ (Genetic engineering medium, RoosterBio) with lentivirus (LV) encoding the SIRPα construct manufactured by Flash Therapeutics. LV volume for transduction was determined by titer and the multiplicity of infectivity (MOI). For LV transduction, the culture medium was replaced with a transduction medium that contained LV and RoosterGEM™, while control MSCs used RoosterGEM^™^ without LV. After incubation of cells with RoosterGEM^™^ for 24 h, the media was exchanged to RoosterNourish™. No rinses were performed before medium replenishment. The cells were maintained for 24 h, seeded into appropriate vessels, and expanded for 4-5 days until the cell confluency reached 80 %. The cells were harvested, formulated in Cryostor 5 (Biolife Solutions), filled into 2 mL cryovials (Corning), and cryopreserved at passage 2 using a controlled rate freezer and stored in the vapor phase of liquid nitrogen. These vials are considered the SIRP-MSC from which all seed train expansions started. SIRP-MSCs were analyzed for expansion ability, MSC characteristics, MSC markers, and SIRPα expression. Mesenchymal stem/stromal cells (MSCs) used in this study were originally derived from human bone marrow aspirate obtained from healthy consenting adult donors from within the United States.

### 3D bioreactor-based upstream process for EV condition media generation

SIRP-MSCs were thawed and expanded in CellBIND^®^ Polystyrene CellSTACK^®^ Chamber (Corning) for 2 passages prior to bioreactor expansion. For generation of the seed train, each passage was considered for expansion of 4 or 5 days to reach a confluency of 80-95 %. Optimization studies were previously conducted to establish the critical process parameters for scaling up the process^56–58^. SIRP-MSCs from the seed train and microcarriers were inoculated into the Ambr^®^250 vessels (Sartorius) or STR^®^50 bioreactor vessels (Sartorius). After inoculation of the Ambr®250 bioreactor, the temperature was maintained at 37 °C, dissolved oxygen (DO) at 100 %, and air flow rate at 24 mL/min with 5 % CO2 in the overlay. In the STR^®^50 bioreactor, the temperature was set to 37 °C, DO control to 50 %, and air flow rate to 1.5 L/min with 5 % CO_2_ in the headspace. RoosterNourish™ was supplemented with a RoosterReplenish™ medium feed (RoosterBio) on day 3 of expansion, per manufacturer instructions. The cells were expanded for 5 days. At the end of the expansion phase, the medium was removed, the cells/microcarriers were washed twice, and then a protein-free, chemically defined low particulate medium, RoosterCollect™-EV (RoosterBio) was added (250 ml final volume in Ambr, 15 L or 50 L in STR50) to the bioreactors to collect EVs at the end of the 5- day culture period. Cell counts and metabolites were measured daily during the expansion phase to monitor cell health. The particle count (particles/mL) was obtained regularly from the bioreactors. Macroscopic and microscopic images of the cells/microcarriers suspension were obtained throughout the process. At the end of the EV collection phase, the bioreactor impellers were stopped and microcarriers were allowed to settle to the bottom of the vessel. After settling, the conditioned media containing EVs was collected through pipetting in the Ambr250 or pumped via the dip tube in the STR50 bioreactor single-use disposable bag. Particle count (particles/mL), particle size and distribution, and SIRPα content were measured. For 2D control flasks served as controls for cell expansion and EV collection, transduced MSCs were seeded into CellBIND^®^ flasks. Cells were cultured for 5 days in an incubator at 37 °C and 5 % CO_2_. At the end of the expansion, the media was aspirated, and the adherent cells were washed before being exchanged into RoosterCollect-EV media for EV collection. TrypLE™ (ThermoFisher) was used when needed for cell harvest from flasks or microcarriers.

### Downstream purification of EV

SIRP-EVs were purified from conditioned medium collected from bioreactors utilizing technologies that are scalable, GMP compatible, and consistent with equipment found at traditional biopharma CDMOs. The conditioned media was treated with Agent V (RoosterBio) per the manufacturer’s instructions prior to clarification. Agent V treated conditioned media was clarified to remove cell debris by pumping through a depth filter (Sartorius). Pre-filter pressure was continuously monitored and maintained per the manufacturer’s recommendations. The clarified medium was concentrated up to 10-fold by volume by filtering through tangential flow filtration (TFF) (Repligen). After concentration, buffer exchange was performed via continuous diafiltration to prepare the concentrated EV solution for chromatography. The concentrated solution was further purified with an AKTA Avant150 chromatography system (Cytiva) using multimodal chromatography resin (Cytiva). The system flow rate, pressures, conductivity, and column UV were controlled and monitored throughout the runs. SIRP-EVs were collected in the flow through fractions, which was consistent throughout scales. The resulting purified EVs were formulated in a sucrose-containing salt buffer via a second TFF diafiltration step (Repligen), then filter sterilized via 0.2 µm membrane (Sartorius) filtration. The resultant purified EV solution was filled into particle-free cryovials (NEST) and stored at −80 °C. To confirm the removal of impurities during the downstream process, intermediate products were analyzed, with total protein measured by Bradford assay (Thermo Fisher), double-stranded DNA (dsDNA) measured by PicoGreen dsDNA assay, and albumin measured using the human albumin ELISA kit (Abcam).

### EV characterization

The size distribution of EVs was analyzed using Dynamic Light Scattering (DLS) with a Zetasizer Nano S90 (Malvern Panalytical). The PDI values were calculated using the Malvern Zetasizer software. The EV particle counts, and zeta potential (ZP) were measured with Nanoparticle Tracking Analysis (NTA) with ZetaView system (Particle Metrix). Data analysis was conducted using ZetaView software version 8.05.16 SP7. Prior to measuring the size distribution and ZP of the EV samples, auto-alignment was completed using a 100 nm polystyrene beads standard solution (WithInstrument, 700074). Before loading the samples, it was ensured that the number of detected particles was below 10, followed by a washing process to commence sample analysis. The background buffer for measurement of size distribution was sample buffer, and deionized water (DW) was used for ZP background buffer. To explore the structure of EVs, Transmission electron microscopy imaging was obtained using the cryo-TEM (FEI Tecnai F20 G2) equipment, and the image analysis was conducted. To quantify protein amount in EVs, BCA assay or Bradford assay was applied, and the results were converted into EV purity (particle/mg). To confirm the percentage of lipid-bound content, MemGlow™ dye was applied and optimized to a working concentration. The dyed samples were evaluated with NanoFCM, and stained events were used to calculate the ratio of bound lipid.

### Liver tissue single cell dissociation

To analyze CD47 expression, necroptotic hepatocyte, cell type-specific biodistribution of EVs, and CD11b^+^ cell depletion, primary hepatocytes and hepatic non-parenchymal cells (NPCs) were isolated using the gentleMACS™ Octo Dissociator with Heaters (Miltenyi Biotec, 130- 096-427) with gentleMACS™ Perfusers (Miltenyi Biotec, 130-128-151), gentleMACS™ Perfusion Sleeves (Miltenyi Biotec, 130-128-752), and Liver Perfusion Kit (Miltenyi Biotec, 130-128-030). For the liver immune cell flow cytometry analysis, NPCs were isolated from HBSS-perfused livers. The livers were transferred to gentleMACS™ C Tubes (Miltenyi Biotec, 130-096-334) and dissociated into single-cell suspension with DMEM containing DNase (Roche, 10104159001) and collagenase (Sigma, C5138) at 37 ℃ utilizing gentleMACS™ Octo Dissociator with Heaters (Miltenyi Biotec, 130-096-427). Dissociated liver tissues were filtered through 70 μm cell strainers (SPL, 93070). Cell suspensions were centrifuged at 300 g for 5 min at 4 ℃. NPC pellet was resuspended in 5 mL RBC lysis buffer (BioLegend, 420301) and incubated for 5 min at 4 ℃. DPBS (Welgene, LB001-02) was added, and suspensions were centrifuged at 300 g for 5 min in 4 ℃. The prepared NPC pellets were resuspended with DPBS for analysis.

### Multiplex immunohistochemistry

The paraffin blocks of liver tissues fixed with 4% formalin were prepared to 4 μm thick slides. The tissue slides underwent sequential steps for staining, heat-drying, dewaxing, and antigen retrieval. After blocking with antibody diluent/block, the slides were stained with primary antibody [F4/80 (Cell Signaling Technology, 70076), HNF-4α (R&D system, PP-H1415-0C), CD47 (GeneTex, GTX53912), Ki67 (Abcam, ab16667), pMLKL (Affinity Biosciences, AF7420)]. Then, tyramide signal amplification was applied to visualize the antigen. The samples were treated with Bone Epitope Retrieval 1 to remove bound antibodies before staining the next antigen. The process from the blocking to retrieval was repeated to stain each primary antibody. Nuclei were visualized with DAPI. The CD47^+^ necroptotic cells were quantified as a percentage relative to the total cells, and Ki-67^+^ cells were quantified relative to the total area (cell/mm^2^).

### Statistical analysis

GraphPad Prism 10 was utilized for statistical analysis. Two-tailed unpaired Student’s t-tests were used to compare two groups. One-way analysis of variance (ANOVA) with Tukey’s post-hoc test was employed for comparisons between multiple groups, and two-way ANOVA with Sidak’s post-hoc test was used for analyzing multiple parameter samples. The survival rate was analyzed using Kaplan–Meier log-rank analysis. The proteomic analysis data was evaluated by Pearson’s correlation coefficient. Spearman correlation was applied spatial transcriptomic analysis. Heatmap analysis was evaluated by Benjamini-Hochberg adjusted P-values. The values were expressed as the mean ± S.D. for samples. *p < 0.05, ** p < 0.01, ***p < 0.001. Additional information on materials and methods is included in the supplementary information.

## Acknowledgements

This research was supported by SHIFTBIO INC., Korean Fund for Regenerative Medicine funded by Ministry of Science and ICT, and Ministry of Health and Welfare (Grant Number: 23C0111L1), a grant of the Korea Health Technology R&D Project through the Korea Health Industry Development Institute (KHIDI), funded by the Ministry of Health & Welfare, Republic of Korea (Grant Number: RS-2023-KH136648; RS-2023-KH140007), and a grant of the BIG3 Project, funded by the Ministry of SMEs and Startups, Republic of Korea (Grant Number: RS-2022-TI022422). Production of LV was outsourced to Flash Therapeutics (Toulouse, France). Quantifying total RNA amount was outsourced to Macrogen (Seoul, Republic of Korea). Multiplex immunohistochemistry (multiplex IHC) was conducted by outsourcing to PrismCDX Co., Ltd. (Gyeonggi-do, Republic of Korea). The biochemistry, hematology, and coagulation analysis were outsourced to DKKorea (Seoul, Republic of Korea). Cryo-TEM (FEI Tecnai F20 G2) imaging was conducted at Korea Institute of Science and Technology Advanced Analysis Center (Seoul, Republic of Korea). Proteomic profiling was analyzed by outsourcing to Bertis (Gyeonggi-do, Republic of Korea).

## Authors contributions

S.K., G.-H.N., G.B.K., and I.-S.K. conceived the idea. S.K., Y.K.K., G.-H.N., and G.B.K. designed the research. S.-Y.P. designed the genetic constructs and performed the cloning. I.K. T.D.S., J.J., E.Z., and S.L. developed the manufacturing process. S.K., Y.K.K., J.K., M.K., S.P., M.K.J., and Y.C. performed the *in vitro* experiments. S.K., Y.K.K., S.K., Y.-S.C., H.J., Y.C., and G.B.K. the *in vivo* experiments. J.W. supported *in vivo* experiments. S.K., Y.K.K., S.K., I.L., J.K., T.D.S., J.J., E.Z., S.L., H.C., J.P. and G.B.K. analyzed the results. S.K., Y.K.K., I.K., G.B.K. and G.-H.N. wrote the manuscript with input from all authors. G.B.K., G.-H.N., and I.-S.K. co-led the study.

## Data Availability

All relevant data generated in this study are available within the article and its Supplementary Information files, Raw data files or from the corresponding author upon reasonable request. The source data is provided with this paper. The publicly available spatial transcriptomic (ST) dataset has been deposited to the National Center for Biotechnology Information Gene Expression Omnibus (GEO) with the accession number GSE223560. The reference single-cell RNA-seq data specific to APAP-induced liver damage and regeneration can be accessed at Zenodo via the dataset identifier https://zenodo.org/records/6035873. The proteomics RAW data are included in the source data files, and the bulk RNA sequencing data are included in the source data and under accession number GSE281951.

