## Supplementary materials for "Dual-mode action of scalable, high-quality engineered stem cell-derived SIRPα-extracellular vesicles for treating acute liver failure"

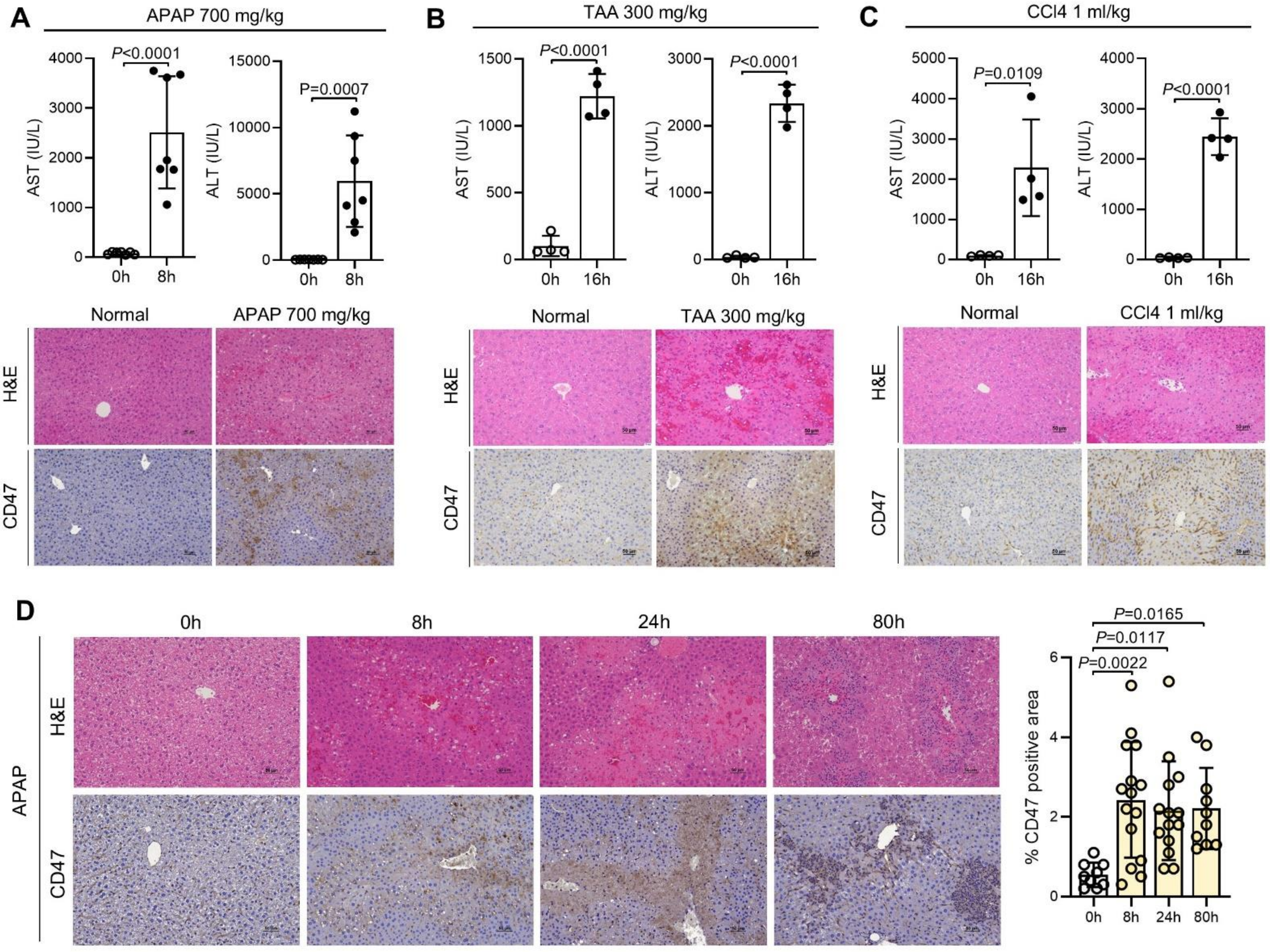

**Supplementary Fig. 1: Assessment of serum AST and ALT levels and histological evaluation in ALF models induced by various agents.** (A-C) Serum AST and ALT levels (upper) and representative images of H&E staining and CD47 IHC analysis from all ALF models (bottom) induced by 700 mg/kg APAP (left, n=7), 300 mg/kg TAA (center, n=4) and 1 ml/kg CCl4 (right, n=4). (D) Representative images of H&E staining and CD47 IHC (left) and quantification of CD47-positive area (right) at 0, 8, 24 and 80 h after ALF induction by APAP. Bar graph data are presented as mean ± S.D. Statistical significance was determined by two-tailed unpaired student's t-test (A-C), and one-way ANOVA with Tukey's post-hoc test (D).

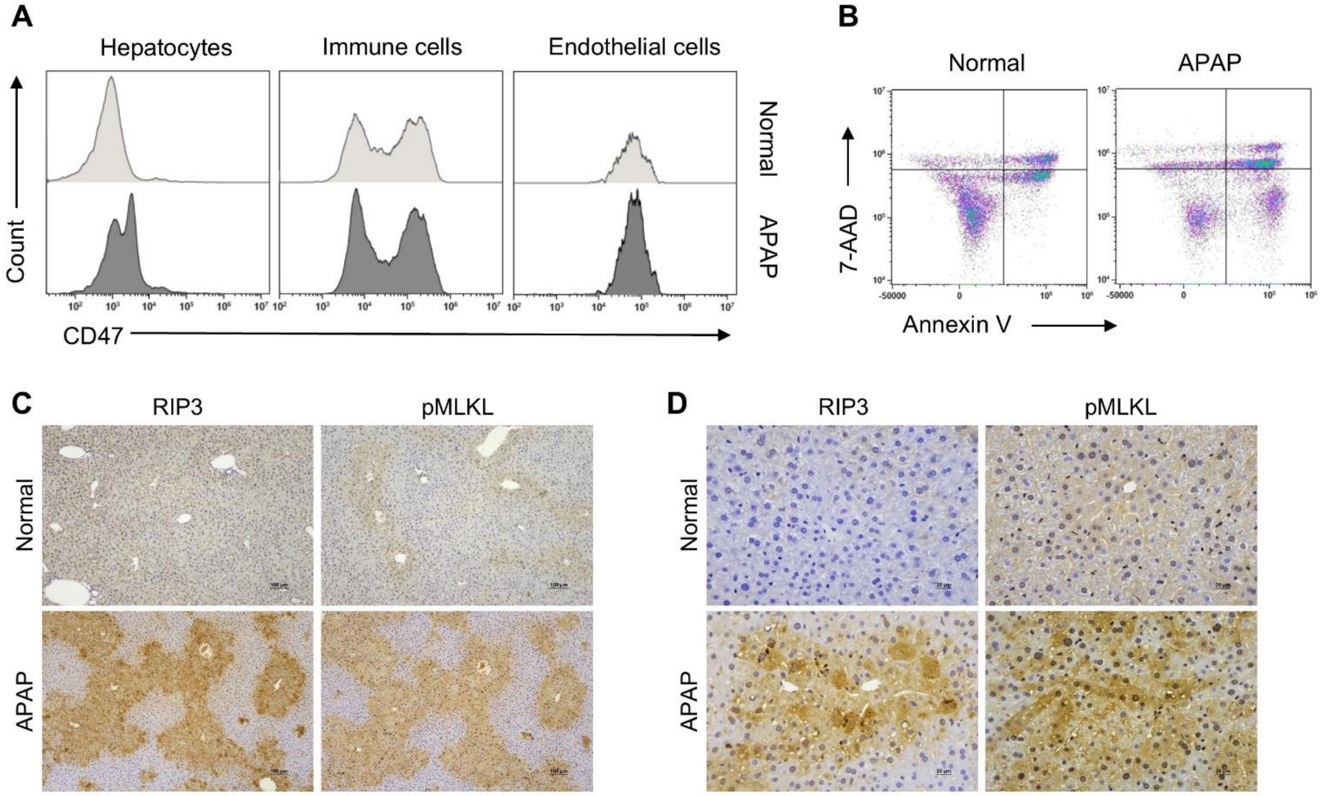

**Supplementary Fig. 2: CD47 is overexpressed on necroptotic hepatocytes in ALF models.** (A) CD47 expression levels of cell populations within liver tissue. The original plot from Fig. 1B. (B) The representative flow cytometry plots of Annexin V/7-AAD analysis to assess early apoptosis, late apoptosis/necroptosis, and necrosis in Normal and APAP-induced ALF mice. The original plot from Fig. 1D. (C and D) Expression of RIP3 and pMLKL in liver samples from normal and APAP- ALF groups at 100X magnification (C) and 400X magnification (D).

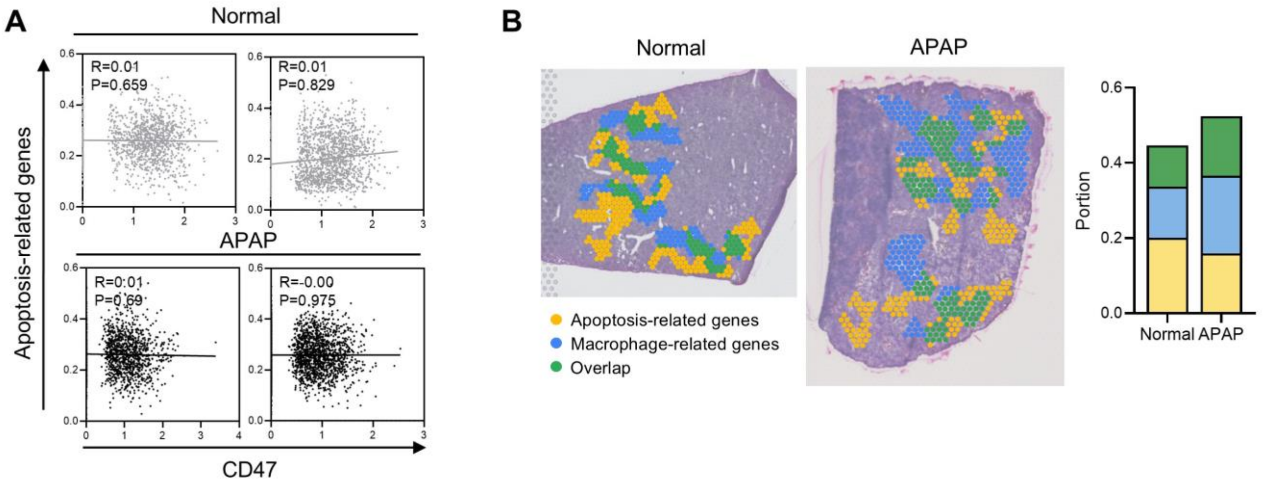

**Supplementary Fig. 3: Spatial transcriptomic analysis of CD47, macrophage and apoptosis-related genes in APAP-ALF.** (A) Scatter plot showing the Spearman correlation (R) between apoptosis-related gene scores and CD47 RNA levels in normal and APAP-ALF liver tissues. (B) STopover analysis illustrating the spatial overlap and interactions between macrophage-related and apoptosis-related gene regions in normal and APAP-ALF liver tissues. Apoptosis-related regions are marked in yellow, macrophage-related regions in blue, and overlapping regions in green. The right panel quantifies the proportion of each region.

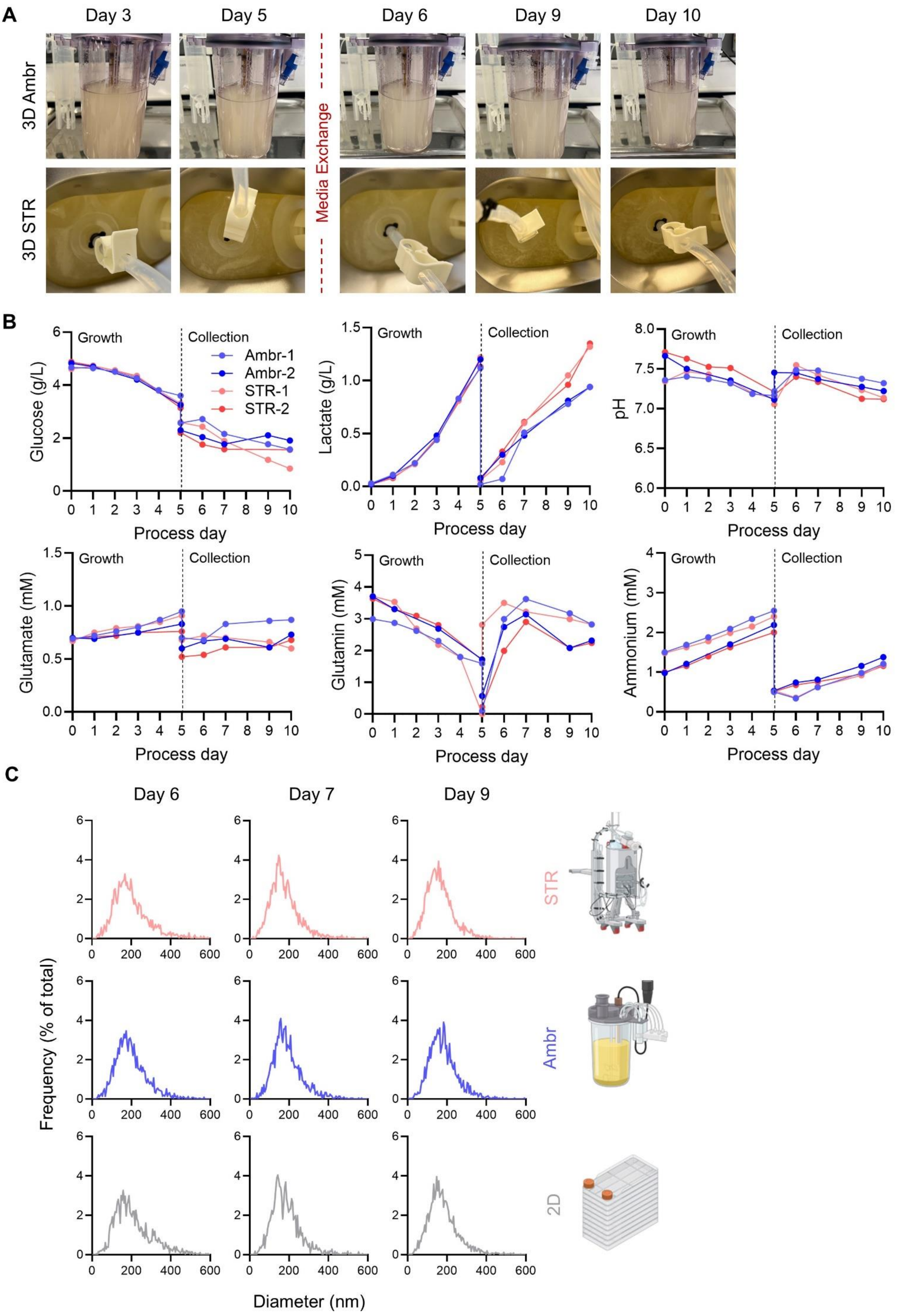
A
Day 3
Day 5
Day 6
Day 9
Day 10
3D Ambr
3D STR
Media Exchange
B
Growth
Collection
Ambr-1
Ambr-2
STR-1
STR-2
Glucose (g/L)
Lactate (g/L)
pH
Glutamate (mM)
Glutamin (mM)
Ammonium (mM)
Process day
C
Day 6
Day 7
Day 9
STR
Ambr
2D
Frequency (% of total)
Diameter (nm)

**Supplementary Fig. 4: Comparison of SIRP-MSC growth in 3D bioreactor systems and 2D control across different scales.** (A) Time course images tracking consistent suspension in STR and Ambr bioreactor vessels. (B) Glucose, lactate, pH, glutamate, glutamine, and ammonium profiles during 15 L Run 1 (STR-1), the Ambr250 satellite from 15 L Run 1 (Ambr-1), 15 L Run 2 (STR-2), and the Ambr250 satellite from 15 L Run 2 (Ambr-2). (C) Particle size distributions measured by NTA at different time points during the collection phase across three tested platforms: 3D STR, 3D Ambr, and 2D culture.

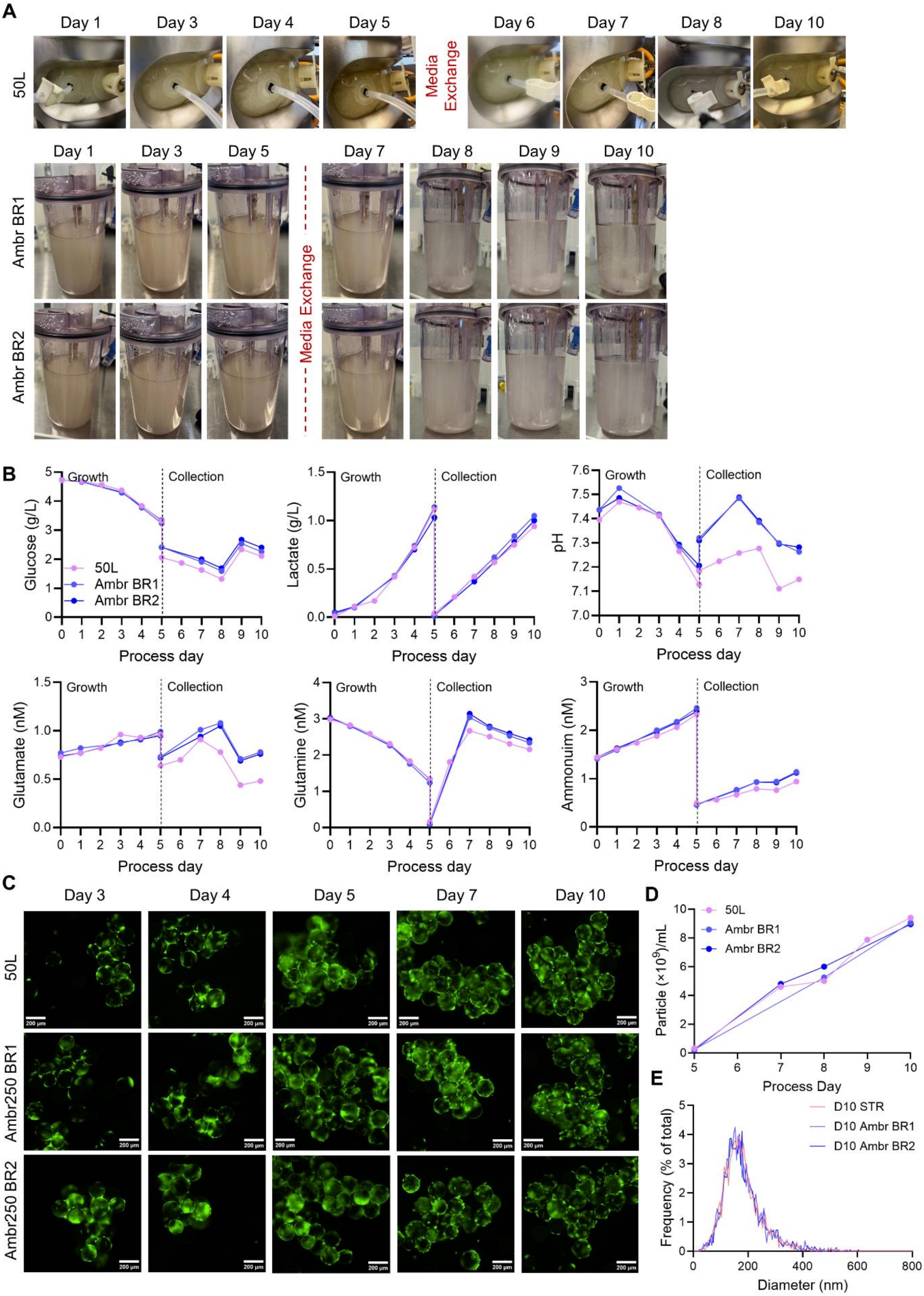
A
Day 1
Day 3
Day 4
Day 5
Day 6
Day 7
Day 8
Day 10
50L
Media Exchange
Day 9
Ambr BR1
Ambr BR2
B
Growth
Collection
Glucose (g/L)
Lactate (g/L)
pH
Glutamate (nM)
Glutamine (nM)
Ammonuim (nM)
Process day
50L
Ambr BR1
Ambr BR2
C
Ambr250 BR1
Ambr250 BR2
D
Particle (×10^9)/mL
Process Day
E
D10 STR
D10 Ambr BR1
D10 Ambr BR2
Frequency (% of total)
Diameter (nm)

**Supplementary Fig. 5: Scale-up in a 3D bioreactor validated by comprehensive analyses across 50 L STR and 250 ml Ambr system.** (A) Observation of macroscopic cell growth over time in bioreactors and Ambr250 control vessels. (B) Metabolic profiles, including glucose, lactate, pH, glutamate, glutamine, and ammonium, during the 50 L STR operation, the Ambr250 vessel (BR1), and the Ambr250 vessel with sparge (BR2). (C) Cell and microcarrier attachment visualized by green calcein AM staining over the culture period. (D) EV particle production over the collection phase. (E) Comparison of size distribution measured by NTA across different scales.

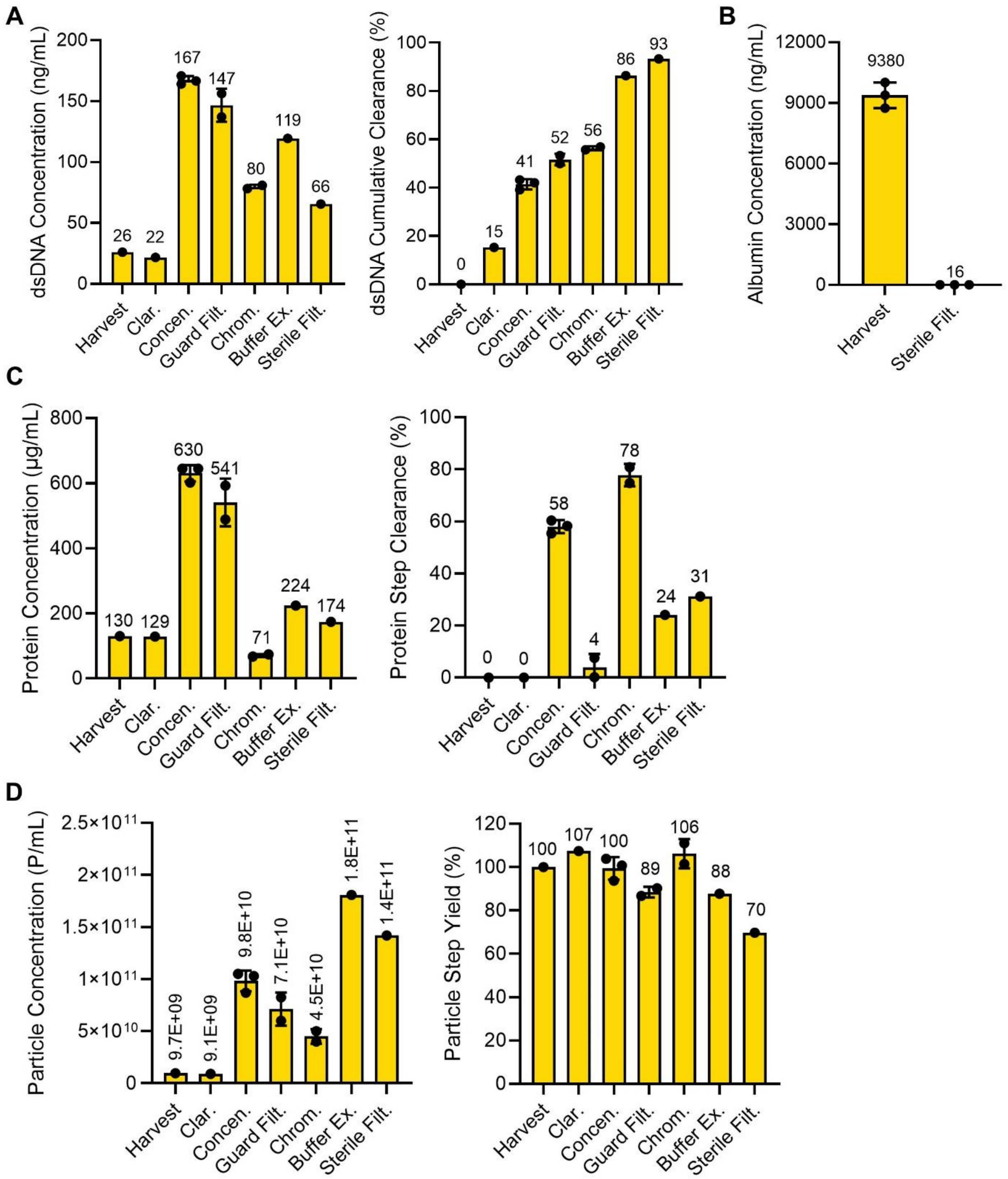

**Supplementary Fig. 6: Comprehensive analyses of downstream process development for SIRP-EV production using 3D cell expansion-based EV isolation.** (A) Quantitation and cumulative clearance of dsDNA in samples. (B) Measurement of human serum albumin concentration in samples. (C) Determination of total protein concentration and step clearance percentage in samples at each processing stage. (D) Assessment of particle concentration and step yield in samples at each processing stage. (A-D) Bar graph data are presented as mean ± S.D. (A, C, D) Bars represent the mean of independent process replicate points using mean analytical test results. (B) Albumin tests included n=3 technical test replicates. The numbers displayed on the bar graph correspond to the exact numerical values of the bar. Clar, clarification; Concen, concentration; Guard Filt, guard filtration; Chrom, chromatography; Buffer Ex, buffer exchange; Sterile Filt, sterile filtration.

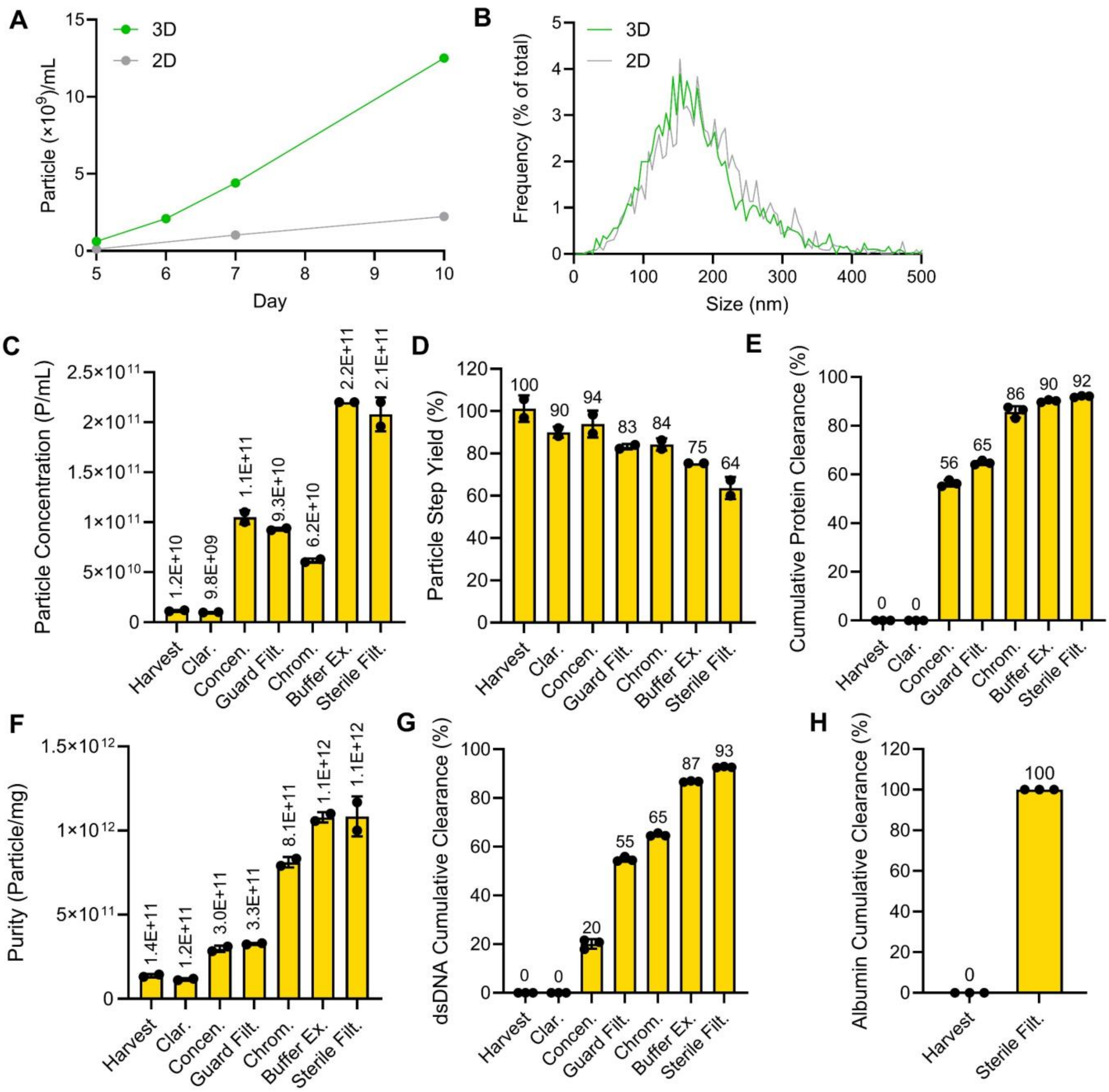

**Supplementary Fig. 7: Overview of upstream and downstream processes of C-EV.** (A) The particle number of EV during production over the collection phase from 3D and 2D culture systems. (B) Size distributions measured by NTA across different culture systems. (C) EV particle concentration in each downstream process. (D) Achieved cumulative yield in the downstream process. (E) Total protein quantitation. (F) Purity of particles per mg of protein. (G) dsDNA quantitation. (H) Quantitation of human serum albumin. (C-H) Bar graph data are presented as mean ± S.D., with bars representing the mean of analytical test replicate data points. The numbers displayed on a bar graph represent the exact numerical values of the bar. Clar, clarification; Concen, concentration; Guard Filt, guard filtration; Chrom, chromatography;

Buffer Ex, buffer exchange; Sterile Filt; sterile filtration

**A**

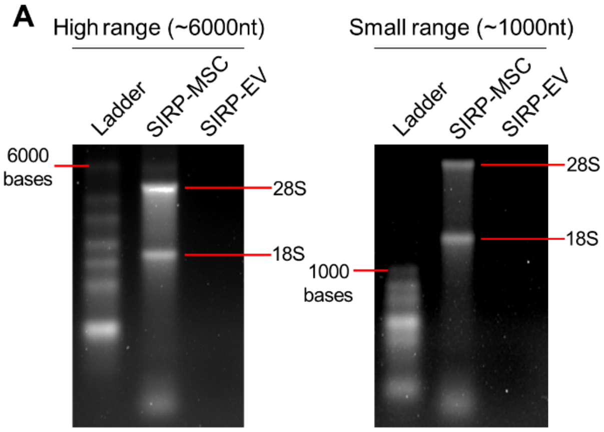

**B**

| | Sample | RNA conc. (ng/µl) | RNA amount per 1E+9 EV (ng) |
|---|---|---|---|
| 2D USP/ Research grade DSP | C-EV | 9.739 | 8.764 |
| | SIRP-EV | 3.616 | 3.254 |
| 3D USP/ GMP grade DSP | C-EV | 1.838 | 1.103 |
| | SIRP-EV | 1.465 | 0.879 |

**Supplementary Fig. 8: Qualitative and quantitative analysis of RNA in SIRP-EVs.** (A) RNA profile of SIRP-MSCs and SIRP-EVs from a 3D bioreactor depicted by electrophoresis with high-range (left) and small-range (right) reference ladders. (B) Measurement of RNA yield from $1\times 10^{11}$ EVs, based on the isolation process, was quantified by RiboGreen assay.

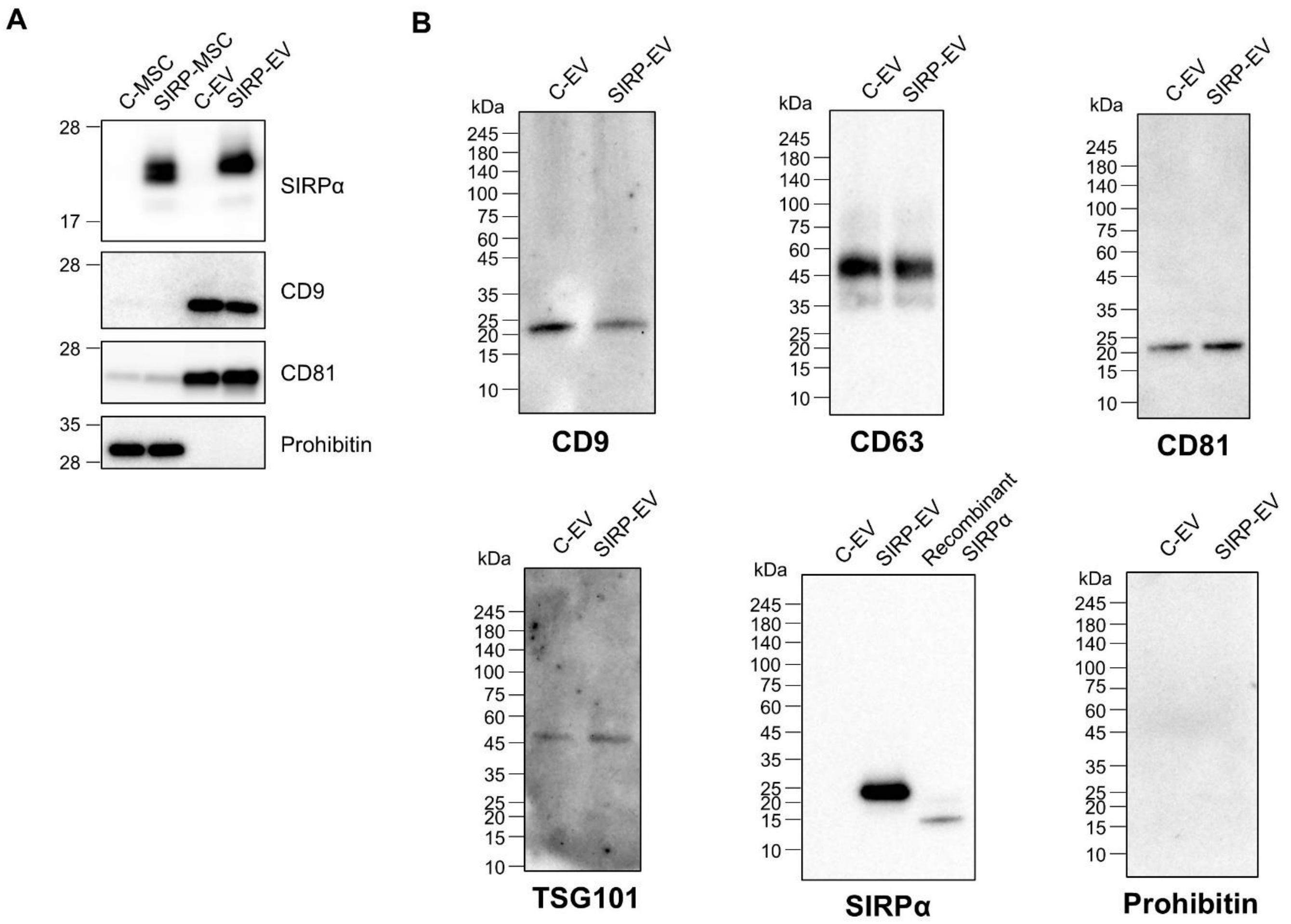

**Supplementary Fig. 9: Characterization of EVs by western blotting.** (A) Western blotting analysis conducted on cell lysates (10 µg) and EVs (2 µg) to examine SIRPα, EV marker proteins (CD81, CD9), and the mitochondrial protein prohibitin. (B) Full scans of the western blots in figure 3 are provided.

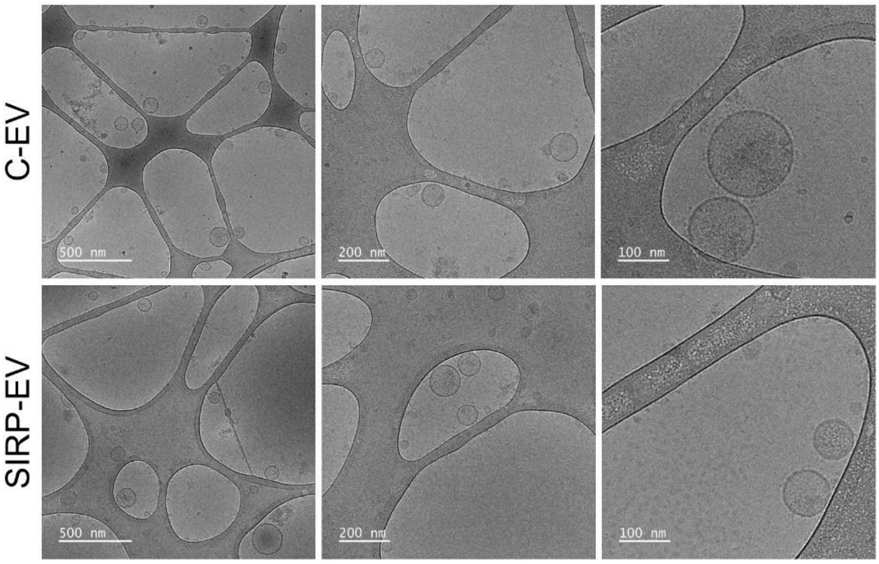

1 **Supplementary Fig. 10: Characterization of EVs by cryo-TEM.** Images captured from multiple fields and at different magnifications. Scale bar, 500 nm, 200 nm, and 100 nm.

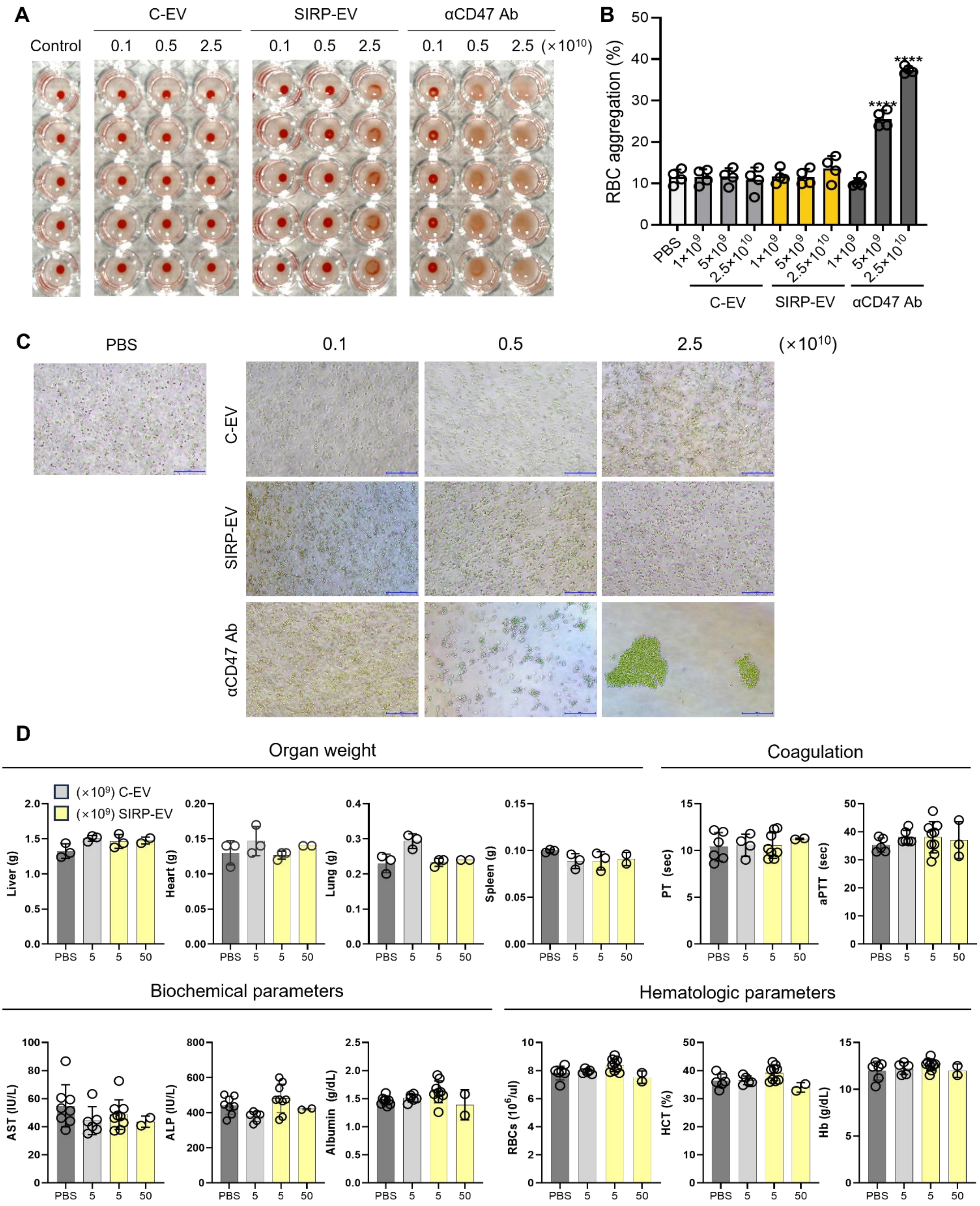

**Supplementary Fig. 11: Assessment of RBC aggregation and toxicity parameters following treatment with EVs and anti-CD47 antibody.** (A-C) Hemagglutination assay using serial five-fold diluted samples of EVs and anti-CD47 antibody. (A) Representative photographs visualizing hemagglutination (B) Quantification of RBC aggregation percentage following treatments, ****P < 0.0001. (C) Representative microscopic images of RBCs. Scale bar, 500 µm. (D) Assessment of organ weight, coagulation profiles, and biochemical and hematologic parameters following administration of C-EVs or SIRP-EVs. Bar graph data are presented as mean ± S.D. Statistical significance was determined by one-way ANOVA with Tukey's post-hoc test. PT, Prothrombin time; aPTT, Activated partial thromboplastin time; AST, Aspartate aminotransferase; ALP, Alkaline phosphatase; RBCs, Red blood cells; HCT, Hematocrit; Hb, Hemoglobin.

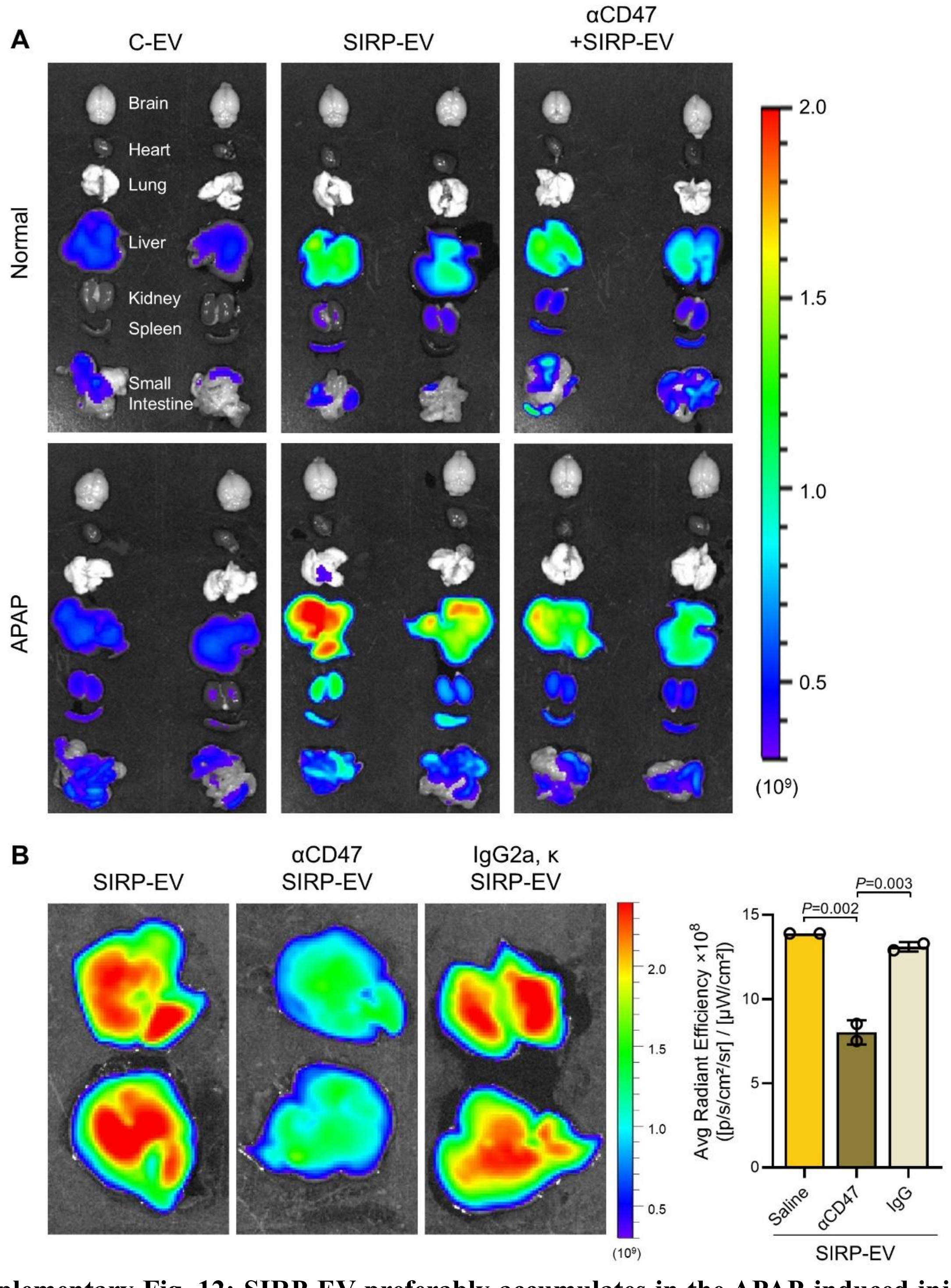

**Supplementary Fig. 12: SIRP-EV preferably accumulates in the APAP-induced injured liver.** (A) *Ex vivo* imaging of the organ distribution in mice 24 h after intravenous administration of 2.5 × $10^{10}$ Cy5.5-labelled SIRP-EVs. (B) *Ex vivo* imaging of liver in mice 24 h after intravenous administration of 2.5 × $10^{10}$ Cy5.5-labelled SIRP-EVs. Mice were either untreated, pre-blocked with CD47 antibody, or pre-blocked with IgG (left). Quantitative fluorescence signals from *ex vivo* livers are presented (right). Bar graph data are presented as mean ± S.D. Statistical significance was determined by one-way ANOVA with Tukey's post-hoc test.

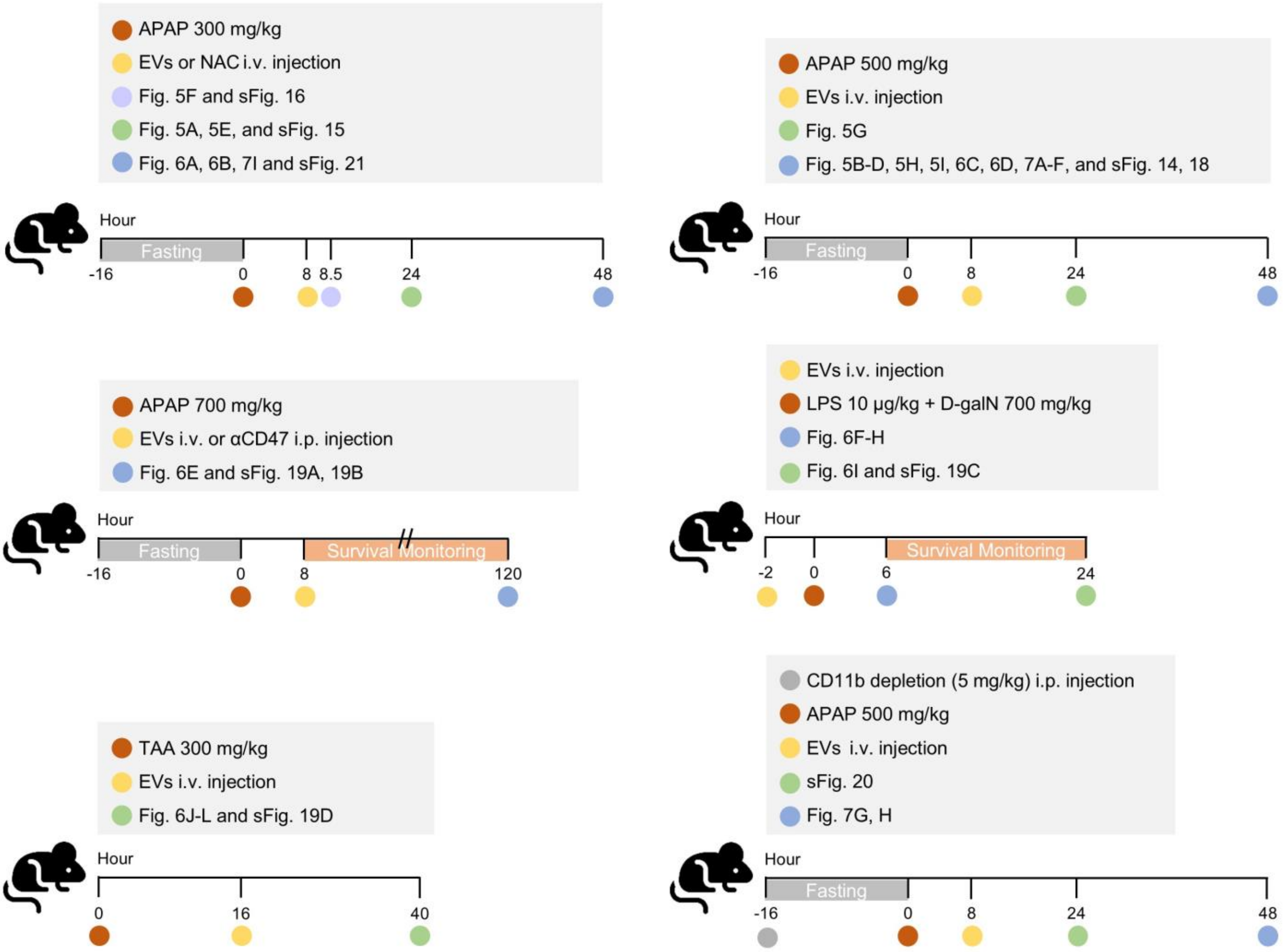

**Supplementary Fig. 13: Schematic schedule of ALF mouse models.** This timeline illustrates the induction conditions for the animal model and therapeutic schedule, including drug administration and injection routes, with related figures.

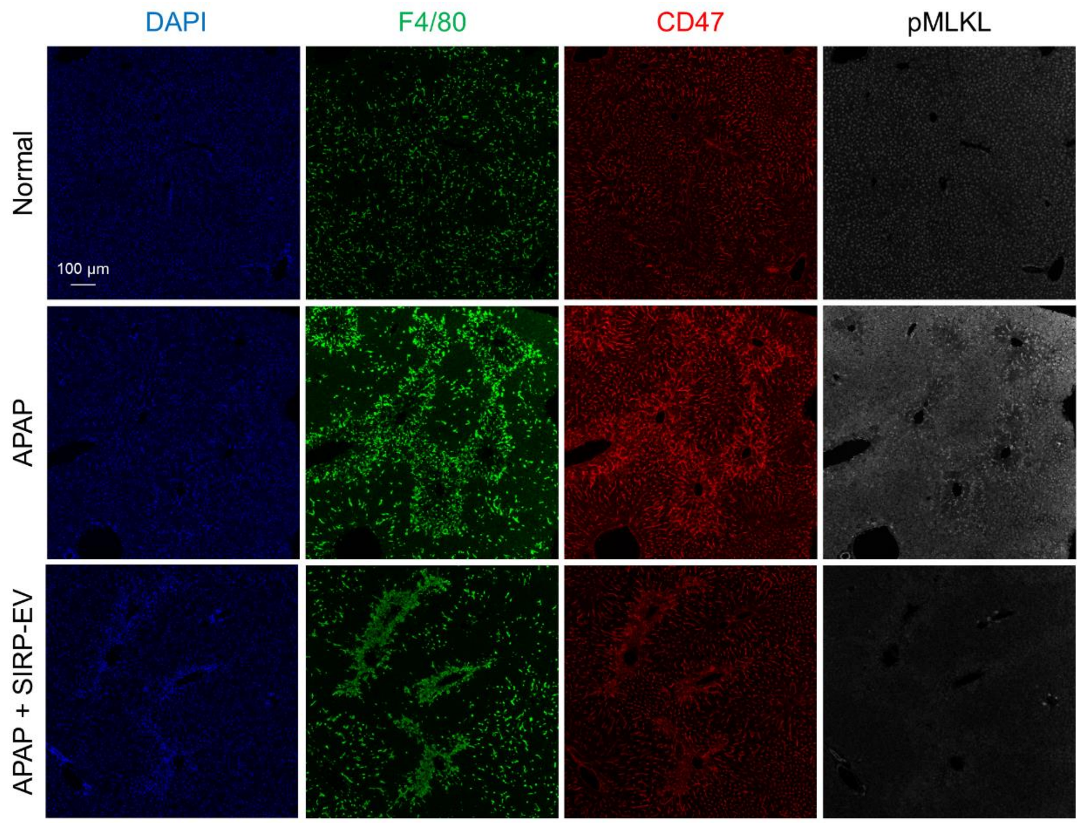

**Supplementary Fig. 14: CD47-positive necroptotic cells in liver tissue were detected using multiplex IHC.** Representative separate staining images depict DAPI (blue), F4/80 (green), CD47 (red), and pMLKL (white). Following the administration of SIRP-EV in APAP-ALF mice, a reduction in CD47-positive necroptotic cells was observed within the liver tissue.

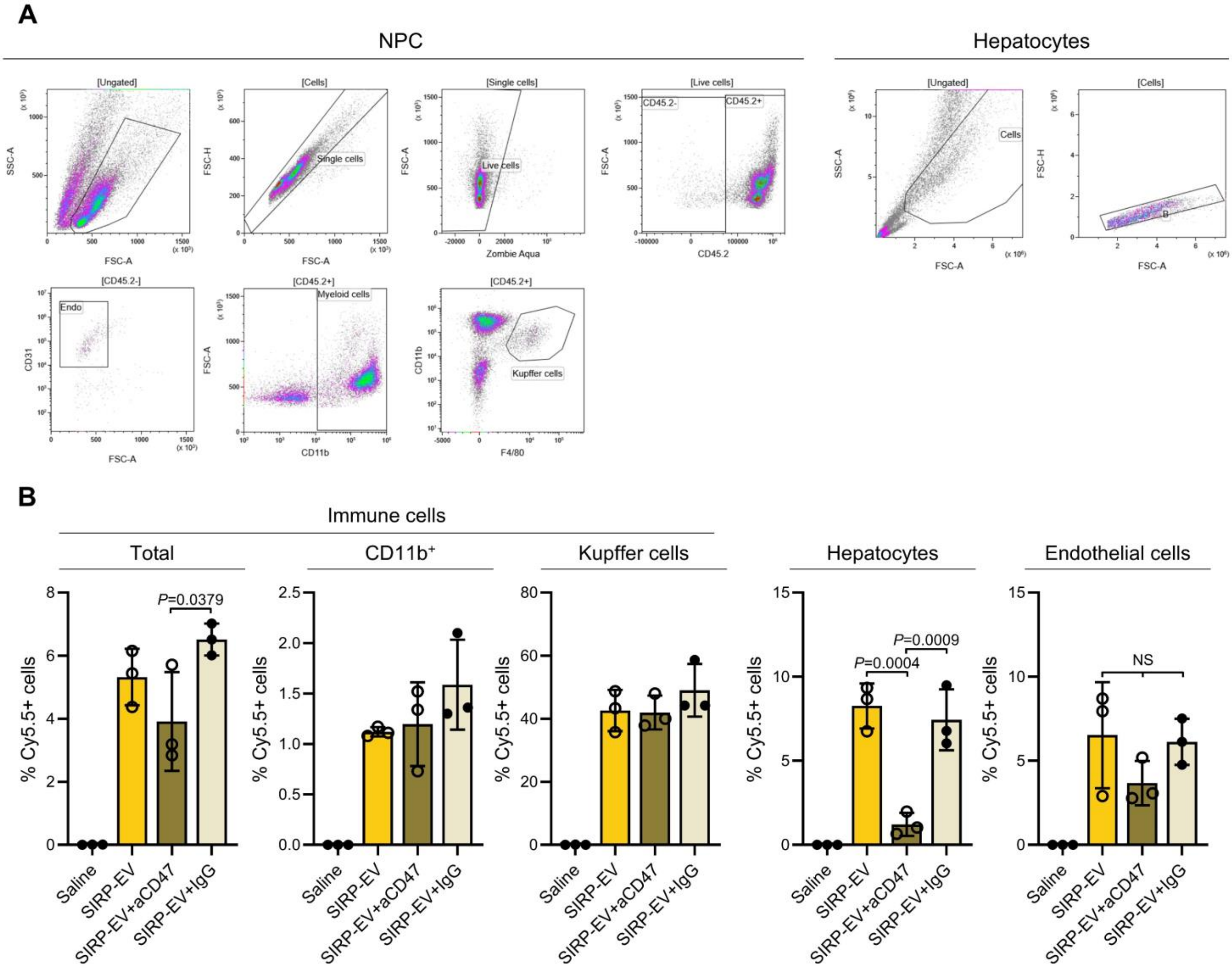

**Supplementary Fig. 15: CD47-dependent hepatocytes specific *ex vivo* biodistribution of SIRP-EVs.** (A) Gating strategies for liver single-cell suspensions to distinguish non-parenchymal cells (NPCs) and parenchymal cells. Specific markers were applied to analyze NPCs, including myeloid cells ($CD45.2^{+}CD11b^{+}$), Kupffer cells ($CD45.2^{+}CD11b^{+}F4/80^{+}$) and endothelial cells ($CD45.2^{-}CD31^{+}$). (B) Cell type-specific biodistribution of SIRP-EV in the CD47-dependent manner in APAP-ALF mice was analyzed using flow cytometry. Mice were either untreated, pre-blocked with CD47 antibody, or pre-blocked with IgG. Bar graph data are presented as mean ± S.D. Statistical significance was determined by one-way ANOVA with Tukey's post-hoc test.

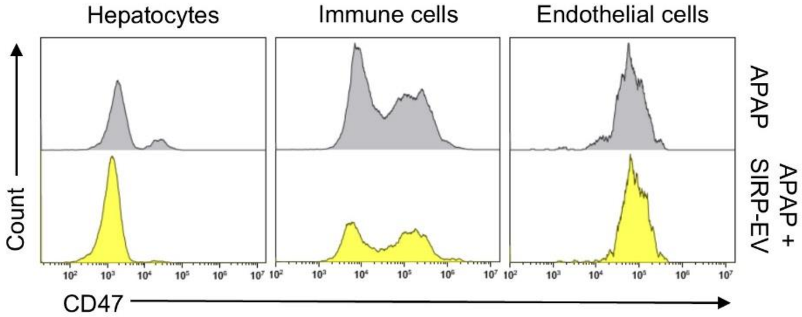

**Supplementary Fig. 16: Changes in CD47 expression in liver tissue induced by SIRP-EV treatment in the APAP-ALF model.** CD47 expression levels of cell populations within liver tissue. The original plot from Fig. 5F.

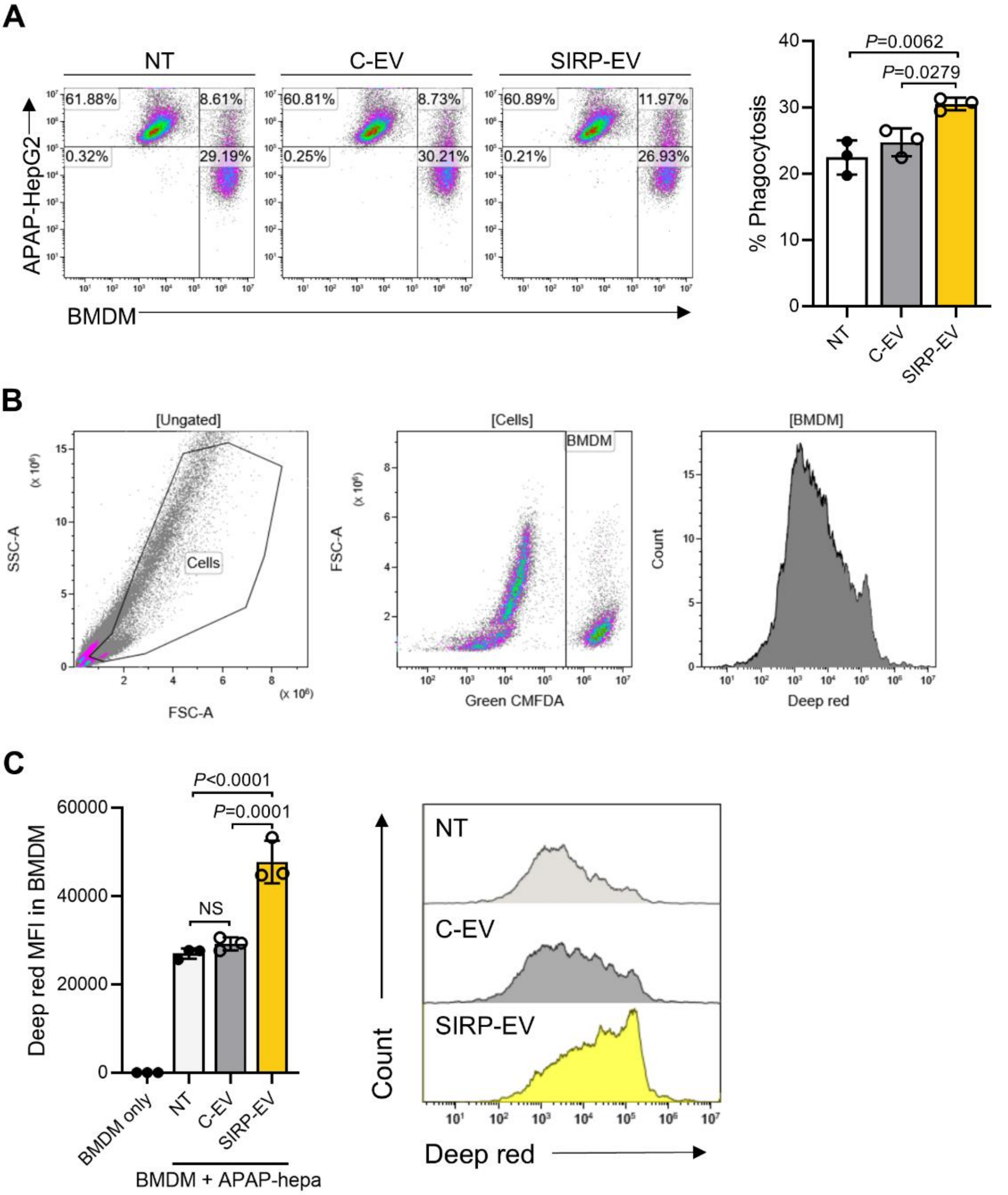

**Supplementary Fig. 17: Flow cytometry analysis of BMDM phagocytosis of APAP-treated HepG2 cells and primary hepatocytes isolated from APAP-ALF models.** (A) Flow cytometry plot showing APAP-treated HepG2 cells stained with Deep Red and BMDMs stained with Green CMFDA (left) Quantification of the percentage of BMDMs containing APAP-treated HepG2 cells is presented on the right (n=3 per group). (B-C) Flow cytometry analysis showing BMDMs labeled with Green CMFDA engulfing hepatocytes, which were isolated from APAP-ALF models and labeled with Deep Red. (B) Gating strategy for identifying BMDMs co-cultured with hepatocytes. (C) Quantification of the mean fluorescence intensity (MFI) of Deep Red from hepatocytes in BMDMs (left) (n=3 per group). Representative histogram of Deep Red MFI in hepatocytes in BMDMs is shown on the right. Bar graph data are presented as mean ± S.D. Statistical significance was determined using one-way ANOVA with Tukey's post-hoc test.

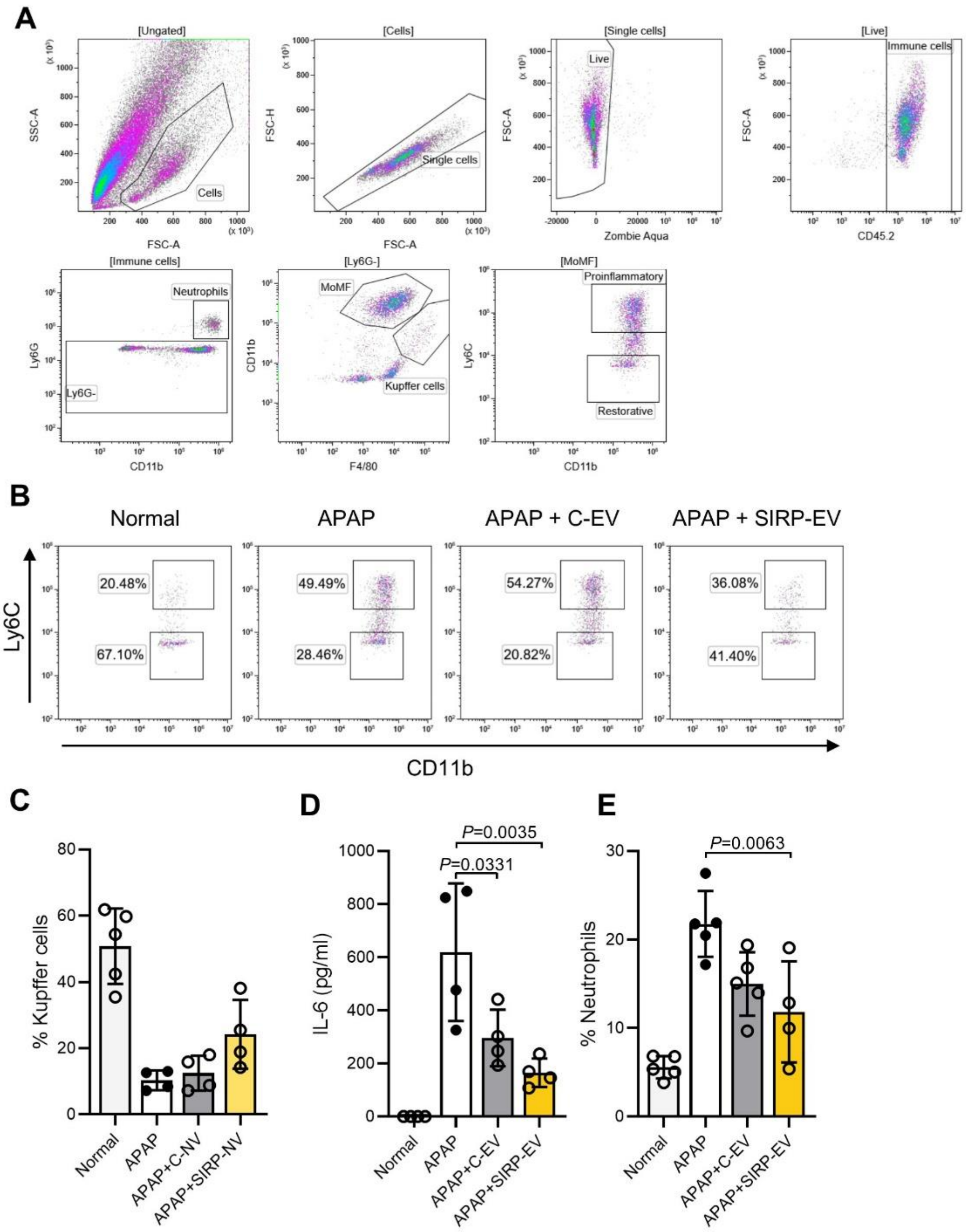

**Supplementary Fig 18. SIRP-EV outperforms C-EV in APAP-ALF models, attributed to SIRPα expression.** (A-D) Therapeutic effects of $4 \times 10^9$ SIRP-EVs in APAP-induced ALF models. (A) Gating strategies for liver single-cell suspensions in APAP-ALF models. (B) Representative flow cytometry images showing Ly6C expression in liver $CD11b^{high}$ $F4/80^{low}$ MoMFs. (C) Kupffer cells as a percentage of total liver $CD45^+$ leukocytes (n=4 or 5 per group). (D) Serum IL-6 levels 48 h post-APAP induction (n=4 per group). (E) Ly6G-positive neutrophils as a percentage of total liver $CD45^+$ leukocytes (n=4 or 5 per group). Bar graph data are presented as mean ± S.D. Statistical significance was determined by one-way ANOVA with Tukey's post-hoc test (C-E).

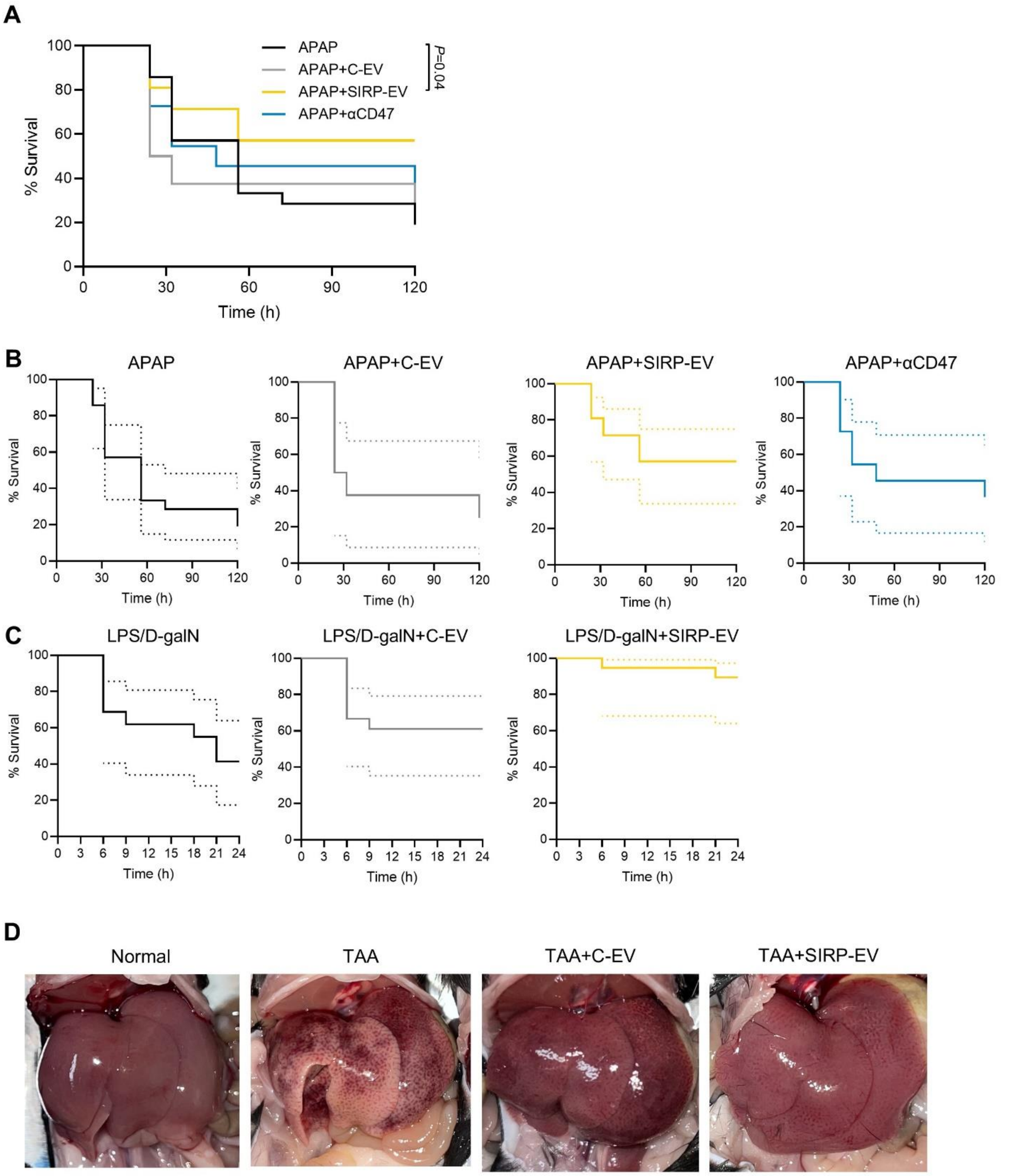

**Supplementary Fig. 19: SIRP-EV demonstrates superior efficacy over C-EV in diverse ALF models.** (A) Kaplan–Meier survival curves for mice with ALF induced by a 700 mg/kg dose of APAP, receiving the indicated treatments (n=8 to 21 per group, three independent experiments). (B and C) 95% confidence intervals for the Kaplan-Meier curves as supplementary figures for both Fig. 6I (B) and Supplementary Fig. 18A (C). (D) Representative gross histology of livers from an ALF model induced by TAA.

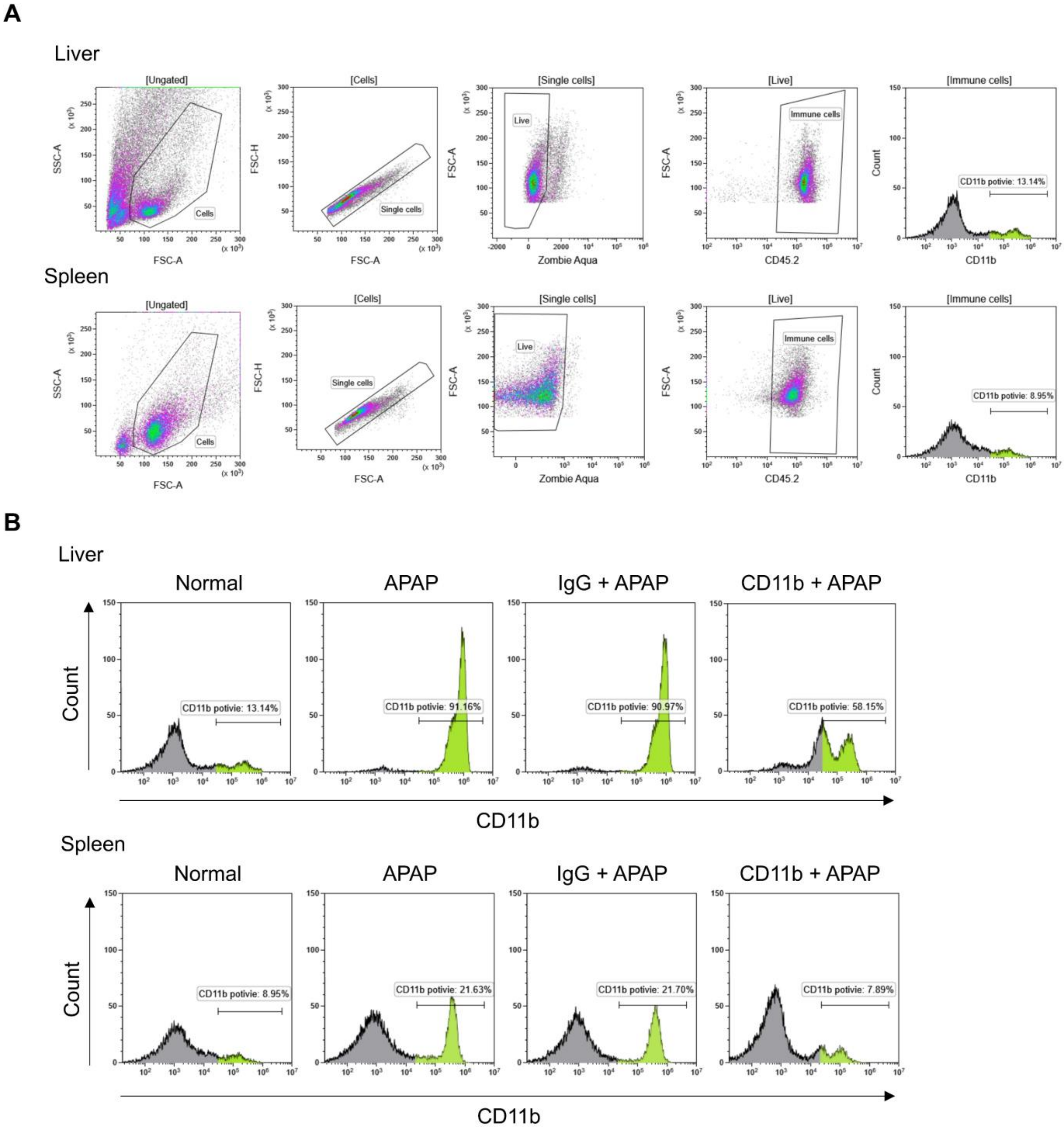

**Supplementary Fig. 20: CD11b$^+$ cell reduction in APAP-ALF model.** (A) Gating strategy for liver and spleen single-cell suspensions to confirm the reduction of CD11b$^+$ cells. (B) Representative histograms from flow cytometry analysis showing depletion levels of CD11b$^+$ cells in livers and spleens (n=4 per group).

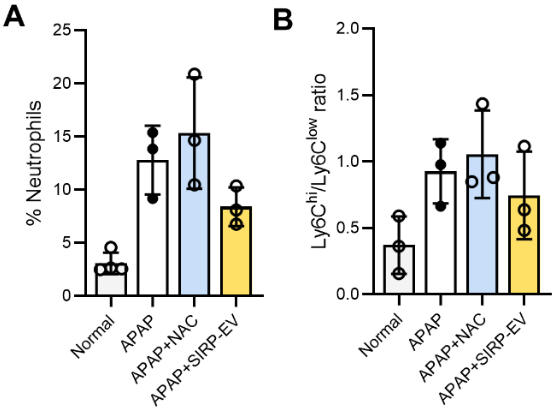

**Supplementary Fig. 21: Competitive advantages of SIRP-EV over N-acetylcysteine in therapeutic application.** (A) Ly6G-positive neutrophils as a percentage of total liver CD45$^{+}$ leukocytes (n=3 per group). (B) Ly6C$^{low}$ to Ly6C$^{high}$ ratio of total liver CD11b$^{high}$ F4/80$^{low}$ MoMFs (n=3 per group). Bar graph data are presented as mean ± S.D. Statistical significance was determined by one-way ANOVA with Tukey's post-hoc test. Only statistically significant comparisons are shown.

| Type | Genes |
| --- | --- |
| Necroptosis related genes | Tnf, Tnfrsf1a, Tradd, Traf2, Traf5, Ripk1, Birc2, Birc3, Xiap, Rbck1, Rnf31, Sharpin, Spata2l, Spata2, Cyld, Fadd, Casp8, Cflar, Ripk3, Cybb, Camk2a, Camk2d, Camk2b, Camk2g, Slc25a4, Slc25a5, Slc25a31, Ppid, Vdac1, Vdac2, Vdac3, Glud1, Glul, Pygl, Pygm, Pygb, Mapk8, Mapk10, Mapk9, Fth1, Pla2g4e, Pla2g4a, Pla2g4b, Pla2g4c, Pla2g4d, Pla2g4f, Alox15, Capn1, Capn2, Smpd1, Mlkl, Pgam5, Dnm1l, Nlrp3, Pycard, Casp1, Il1b, Chmp2a, Chmp2b, Chmp3, Chmp4b, Chmp4c, Chmp6, Vps4b, Vps4a, Chmp1b, Chmp1a, Chmp5, Chmp7, Trpm7, Il1a, Il33, Hmgb1, Tnfsf10, Tnfrsf10b, Fas, Ifna1, Ifna2, Ifna4, Ifna5, Ifna6, Ifna7, Ifna13, Ifna14, Ifna16, Ifnb1, Ifng, Ifnar1, Ifnar2, Ifngr1, Ifngr2, Jak1, Jak2, Jak3, Tyk2, Stat1, Stat2, Stat3, Stat4, Stat5a, Stat5b, Stat6, Irf9, Eif2ak2, Tlr4, Ticam2, Ticam1, Tlr3, Zbp1, Usp21, Sqstm1, Hsp90aa1, Hsp90ab1, Tnfaip3, Parp1, Bid, Bax, Aifm1, H2al3, Ppia, Bcl2 |

**Supplementary Table 1.** Fig. 1G-I shows the list of 121 genes used for scoring necroptosis-related genes.

| Abbreviation | Expanded Form |
|---|---|
| ALF | Acute Liver Failure |
| ALT | Alanine aminotransferase |
| Ambr | Small-scale (250ml) Ambr250 |
| APAP | Acetaminophen |
| APAP-ALF | Acetaminophen induced Acute Liver Failure |
| AST | Aminotransferase |
| BMDM | Bone Marrow Derived Macrophage |
| CCl4 | Carbon Tetrachloride |
| C-EV | Naïve mesenchymal stem cell-derived EV |
| C-MSC | Control Mesenchymal Stem Cell |
| D-galN | D-galactosamine |
| dsDNA | Double stranded DNA |
| ECM | Extracellular Matrix |
| EV | Extracellular Vesicle |
| GOBPs | Gene Ontology Biological Processes |
| H&E | Hematoxylin & Eosin |
| hBM-MSC | Human bone marrow-derived Mesenchymal Stem Cells |
| HEKSIRP-EV | HEK cell-derived SIRP-EV |
| HSA | Human Serum Albumin |
| i.p. | Intraperitoneal |
| IHC | Immunohistochemistry |
| LC-MS/MS | Liquid chromatography with tandem mass spectrometry |
| LPS | Lipopolysaccharide |
| MoMF | Monocyte-derived macrophage |
| MSC | Mesenchymal Stem Cell |
| NAC | N-acetylcysteine |
| RBC | Red Blood Cell |
| SIRP | Signal Regulatory Protein Alpha |
| SIRP-EV | EV expressing SIRPα from genetically MSC |
| SIRP-MSC | Mesenchymal Stem Cells to express SIRPα creating SIRPα-modified MSC |
| ST | Spatial Transcriptomics |
| STR | Large-scale (15L) Biostat STR® |
| TAA | Thioacetamide |
| TFF | Tangential flow filtration |
| ZP | Zeta Potential |

**Supplementary Table 2.** Abbreviation

### Supplementary Materials and Methods

#### Cell culture

The 293FT cells (HEK293 cell derivatives) were maintained in Dulbecco's modified Eagle's medium with high glucose (DMEM, Cytiva, SH30243.01) supplemented with 10 % fetal bovine serum (FBS, Gibco, 12483-020) and 1 % antibiotic-antimycotic (A.A., Gibco, 15240-062). To generate a stable cell line, Plat-E cells were used to produce a retrovirus containing a DNA sequence of interest with a puromycin resistance gene. When the confluency of 293FT cells reached 80-90 %, Plat-E-derived retrovirus transduction was performed. Plat-E cells and 293FT cells were grown at DMEM with 10 % FBS and 1 % antibiotic-antimycotic, and virus-transduced 293FT cells were grown at puromycin-added medium with the same medium of 293FT cell media. The hBM-MSCs (RoosterBio) were maintained with RoosterNourish™ (RoosterBio) in CellBIND® Polystyrene CellSTACK® (Corning) for 2D cell culture. The CD47 KO cell line (Abcam, ab266324) and WT CD47 cell line was purchased from Abcam and maintained following noted manufacturer's handling procedure. To acquire the mouse BMDMs, mouse bone marrow derived primary cells were differentiated and maintained in accordance with the previously reported method[1,2].

#### 2D cell culture-based EV production and purification

The SIRPα transduced 293FT cells and MSCs were utilized to produce EVs ($_{HEK}$SIRP-EV, C-EV, and SIRP-EV). Once the cell confluency reached 90 %, the medium was replaced with serum-free medium composed of DMEM, 1 % GLUTAMAX™ (Gibco, 35050061), and 1 % A.A or RoosterCollect™-EV (RoosterBio). The supernatant was collected after 48 h and subjected to sequential centrifugation at 300 g for 10 min, 2,000 g for 10 min, and 10,000 g for

30 min. The supernatant was then filtered and purified using the KrosFlo® KR2i Tangential Flow Filtration System equipped with hollow fiber filters (TFF, Repligen, Spectrum Labs). The samples were subsequently concentrated and diluted with PBS. Post-TFF, the samples were sterilized by filtration and centrifuged at 150,000 g for 1.5 h to gain pelleted EVs. The EV pellets were resuspended in PBS containing a proteinase inhibitor cocktail (Sigma-Aldrich, 535140).

#### Spatial transcriptomics

##### *Processing of ST Data*

This study used a publicly available spatial transcriptomic (ST) dataset (accession number GSE223560) of APAP-induced liver regeneration, focusing on liver samples from three healthy control mice and two collected 24 hours post-APAP induction. All ST data were processed using the Seurat pipeline (version 4.0.2) in R (version 4.0.5). To integrate multiple datasets, the functions SelectIntegrationFeatures, FindIntegrationAnchors, and IntegrateData were employed. After normalization, spatial features were analyzed by examining associations such as the relationship between necroptosis and CD47 expression by calculating Spearman correlation coefficients.

##### *Necroptosis and Apoptosis Signature Score*

The necroptosis or apoptosis signature scores were calculated to assess its correlation with CD47 expression in the ST data. Using the AddModuleScore function in Seurat, we computed necroptosis gene set scores for each spatial spot. Spearman correlation was then applied to evaluate the association between necroptosis or apoptosis signature scores with CD47 expression across spots. The list of genes was obtained from https://www.genome.jp/entry/pathway. The necroptosis-related gene list is provided in Supplementary Table 1, and the list of apoptosis-related genes is represented below: Aifm1, Apaf1, Bad, Bak1, Bax, Bbc3, Bcl2l11, Bid, Hrk, Pmaip1, Cycs, Casp3, Fas, Fasl, Diablo, Htra2, Endog, Trp53.

***Cell type mapping on ST data***

Spatial cell types were mapped by integrating spatial transcriptomics data with reference single-cell RNA-seq data specific to APAP-induced liver damage and regeneration, available at https://zenodo.org/records/6035873[3]. For this integration, the CellDART algorithm was used[4]. In the mapping process, the number of marker genes for each cell type, as defined by the scRNA-seq data, was set to 30, while other parameters were kept at default to generate cell type maps on the ST data.

***Topological overlap analysis on ST data***

To investigate spatial colocalization patterns between the necroptosis or apoptosis signatures with macrophages in liver tissue, topological overlap analysis was performed using the STopover method[5]. This approach identifies topologically defined regions within the tissue that show local activation or accumulation of specific markers and assesses the extent of overlap between these regions. The proportion of each region relative to the overall tissue was evaluated by counting ST spots in both overlapped and non-overlapped areas. Additionally, SIRPα expression within these regions was analyzed to further characterize cellular interactions.

**RNA analysis**

The total RNAs in cells and EVs were isolated utilizing miRNeasy micro kit (Qiagen, 217084).

To detect various ranges of RNA in samples, RNA electroporation was applied with two standard ladders (Thermo Fisher, SM1831, and SM1821), and the gels were detected by GelDoc Go Imaging System (Bio-Rad). The amount of RNA was quantified using the Quant-it™ RiboGreen RNA assay kit (Invitrogen, R11490), which was applied to RNase-free DNases pre-treated samples, and fluorescence was detected using VICTOR Nivo™ multimode microplate reader (PerkinElmer, HH35000500).

**Flow cytometry**

Hepatic NPCs and primary hepatocytes were isolated from the mouse models following the liver tissue single cell dissociation method. Isolated single NPCs were pre-stained with Zombie Aqua™ (BioLegend, 423102). Then, the cells were pre-incubated with anti-mouse CD16/CD32 (BD Pharminge™, 553142) and stained with fluorescence conjugated antibodies for liver immune cell analysis. Antibodies were purchased from BioLegend [PC7-anti-CD45.2 (109830), APC-anti-CD11b (101212), BV785-anti-CD11b (101243), BV421-anti-CD31 (102424), BV605-anti-Ly6G (127639), FITC-anti-F4/80 (123108) and PB-anti-Ly6C (128014)]. To analyze the cell death phase, primary hepatocytes were stained with FITC-Annexin V and 7-AAD (BioLegend, 640922). The stained cells were analyzed by Beckman CytoFLEX flow cytometry and Kaluza Analysis 2.1.

**Confocal microscopy**

For confocal microscopy, liver tissues extracted from APAP-ALF models were immediately frozen in liquid nitrogen. The frozen tissues were then cryo-sectioned, which were fixed with 4 % paraformaldehyde (Biosesang, P2031-050-00) and underwent a washing process 4 times for 5 min each, followed by blocking in 3 % BSA/PBS solution for 1 h at room temperature. The primary antibodies [pMLKL (Abcam, ab196436) and CD47 (R&D system, AF1866)] were applied to stain slides using a 1 % BSA/PBS solution. After overnight incubation with the primary antibodies, samples underwent a washing process 4 times 5 min each. The secondary antibodies [Anti-rabbit Alexa Fluor 647-conjugated antibody (Jackson ImmunoResearch, AB_2338072) and Anti-goat Alexa Fluor-488-conjugated antibody (Jackson ImmunoResearch, AB_2340439)] were then applied using 1 % BSA/PBS solution. After the washing process, the samples were mounted with DAPI Fluoromount-G solution (SouthernBiotech, 0100-20). The microscopic images were gained with confocal microscopy (Zeiss, LSM800).

**Western blot**

To analyze the expression of CD47, SIRPα, EV markers (CD9, CD63, CD81, and TSG101), and prohibitin, a western blot assay was employed. Protein concentrations of each sample were quantified using the BCA protein assay to ensure the loading of identical amounts of protein. Samples were lysed with RIPA buffer (Cell Signaling Technology) and mixed with Laemmli sample buffer (GenDEPOT). Initially, gels were run at 70 V for 10 min, followed by 100 V for 90 min to 2 h. Proteins were transferred to a PVDF membrane at 110 V for 90 min. After transferring proteins, membranes were blocked with a 5 % skim milk solution or 5 % BSA solution in 1× TBST for 1 h at room temperature. Then membranes were incubated with the primary antibody [CD47 (R&D System, AF1866), β-actin (R&D System, MAB8929), SIRPα (inhouse development), CD9 (System Biosciences, EXOAB-CD9A-1), CD63 (Abcam, ab68418), CD81 (Santa Cruz Biotechnology, sc-166029), TSG101 (Abcam, ab125011), and prohibitin (Novus Biologicals, H00005245-M01)] in a 5 % skim milk solution or 3 % BSA overnight at 4 °C on a low-speed setting. Following primary antibody incubation, membranes were washed with 1× TBST, changing the wash buffer every 10 min, for 1 h at room temperature on a high-speed setting. Then membranes were incubated with the secondary antibody [horseradish peroxidase (HRP)-conjugated anti-mouse IgG (Abcam, ab6728), anti-rabbit IgG (Abcam, ab6721) and anti-goat IgG (Thermo Fisher Scientific, 31402)] in a 5% skim milk solution for 1 h at room temperature on a low-speed setting. After incubation with secondary antibodies, the membrane was processed with enhanced chemiluminescence (ECL) western blotting substrate (Thermo Fisher Scientific, MA, USA), and the image was gained with the Amersham Image Quant™ 800 system.

**Proteomic profiling of EVs**

For the transcriptomic analysis of C-EV, $_{\text{HEK}}$SIRP-EV and SIRP-EV samples, Tandem Mass Tag (TMT) labeling, an isobaric tag reagent, was employed followed by LC-MS/MS analysis. The identified proteins and their quantitative values in each batch were used to compare differences between batches. The number of proteins analyzed for each sample type was 7,391 for both C-EV and SIRP-EV. For sample lysis, EV samples were added chilled acetone with four volume of samples and incubated at -80 °C for 90 min. The reacted samples were centrifuged, removed supernatant, and dried. The dried samples were resuspended in 8 M urea in 100 mM ammonium bicarbonate and subjected to sonication for 5 min to lyse the precipitated samples. After centrifugation at 16,000 rpm for 5 min, the supernatant was transferred to a new tube, and the protein concentration was measured using a BCA assay kit to adjust the same amount of protein to be analyzed. For protein reduction and alkylation, dithiothreitol (DTT) was added to the EV lysates at a final concentration of 10 mM, and the mixture was incubated at 37 °C with shaking at 450 rpm for 30 min to reduce the proteins. After reduction, IAA (iodoacetamide) was added at a final concentration of 25 mM, and the mixture was incubated in the dark at room temperature for 30 min for alkylation. For protein digestion, the alkylated protein samples were diluted with 100 mM ABC (ammonium bicarbonate) to decrease the urea concentration to below 1 M. Trypsin digestion was performed at a 1:25 protein:enzyme ratio, and the reaction was allowed to proceed for 16 h at 37 °C. The trypsin reaction was quenched by adding TFA (trifluoroacetic acid) to a final concentration of 1 % TFA. The prepared samples were TMT labelled, and TMT labelled 100 μg of samples were fractionated into 12 fractions to be analyzed by Nano LC-MS/MS system (Thermo Dionex Ultimate 3000 coupled with Thermo Orbitrap Exploris 480). The mass spectrometric data were matched with human protein sequence provided by UniProt database and analyzed with SAGE software and Gene Ontology Biological Processes (GOBPs) among the top 506 SIRP-EV proteins was established through FunRich software v.3.1.3.

**Bulk RNA-sequencing analysis**

To compare the RNA expression, the APAP (500 mg/kg) induced mice were treated with $_{HEK}$SIRP-EV or SIRP-EV, and the livers were harvested 24 hours after APAP induction. The harvested livers were dissociated into cells following liver tissue single-cell dissociation protocol, and CD11b-positive cells were isolated using CD11b MicroBeads UltraPure (Miltenyi Biotec, 130-126-725). Total RNA from CD11b-positive cells was prepared using Trizol reagent (Invitrogen, 15596026). The quality of RNA (RNA integrity) was assessed using an Agilent Technologies 2100 Bioanalyzer. The size of the RNA library was determined using an Agilent Technologies 2100 Bioanalyzer with a DNA 1000 chip, and the RNA library was quantified following the Illumina qPCR Quantification Protocol Guide. To prepare aligned reads, the trimmed RNA reads were mapped to the reference genome using the HISAT2 program, and then the aligned reads were processed for transcript assembly with the StringTie program. Gene expression profiles were calculated by the read count, normalized FPKM (Fragments Per Kilobase of transcript per Million mapped reads)/RPKM (Reads Per Kilobase of transcript per Million mapped reads), and TPM (Transcripts Per Kilobase Million). To compare different gene expressions among the groups, the significant genes were identified by applying adjusted p-value with Benjamini-Hochberg procedure and categorized into the five gene clusters (Cellular viability, Detoxification/Antioxidation, ECM Remodeling, Immune Modulation, and Vascularization and Perfusion). The list of 404 genes (Source Data Fig. 7F) was applied for DEG (Differentially Expressed Genes) analysis. For the DESeq2 analysis, the negative binomial Wald Test (nbinomWaldTest) was conducted with RLE (Relative Log Expression) normalized counts.

**Cell binding assay**

For SIRPα *in vitro* cell binding analysis, each set of EVs was labeled with sulfo-Cyanine5.5 NHS ester (Lumiprobe, 27320) at a ratio of $5 \times 10^{10}$ particles per 25 ng of Cy5.5 dyes and incubated at 4 °C overnight. The Zeba™ Spin Desalting Columns (ThermoFisher, A57762) were employed to purify the labeled EVs and eliminate free Cy5.5 dyes. The fluorescence intensity of Cy5.5-NHS ester conjugated EVs was analyzed using a microplate reader (PerkinElmer, VICTOR Nivo™) to compare the fluorescence intensity among samples. $2 \times 10^5$ HEK CD47 WT (wild type) or KO (knockout) cells were mixed with 200 μL of 3 % BSA and incubated at 4 °C for 15 min. The samples were then centrifuged at 450 g for 3 min at 4 °C, and the supernatants were discarded. The pellets were resuspended in 200 μL of the desired concentrations of C-EV or SIRP-EV ($5 \times 10^8$, $1 \times 10^9$, or $2.5 \times 10^9$) and incubated at 4 °C for 15 min. Following another centrifugation at 450 g for 3 min at 4 °C, the supernatants were discarded. The pellets were washed with 200 μL of 1 % BSA in PBS and centrifuged at 450 g for 3 min. The final pellets were resuspended in 200 μL of PBS and analyzed by flow cytometry.

##### *In vitro* phagocytosis assay

The fully differentiated BMDMs were labeled with 1 μM CellTracker Green CMFDA (Invitrogen, C7025) for 30 min at room temperature in the dark and the excess dye was removed by following centrifugation. The stained BMDMs were subsequently seeded 24 h before the assay. HepG2 cells treated with 40 μM APAP or primary hepatocytes isolated from 300 mg/kg APAP-ALF model mice were labeled with 0.5 μM CellTracker Deep Red (Invitrogen, C34565). BMDMs and either HepG2 cells or primary hepatocytes were co-cultured in the presence of C-EVs or SIRP-EVs at a cell ratio of 1:2 at 37 °C. After the incubation, all cells were collected, washed, and analyzed using flow cytometry. Phagocytosis levels were quantified by calculating the percentage of HepG2 cells (Deep Red$^+$ cells) within the total BMDM population (Green CMFDA$^+$ cells). For analysis of phagocytic activity, the MFI of Deep Red in primary hepatocytes internalized by BMDMs was quantified.

##### CD11b$^+$ myeloid cell reduction in the APAP induced ALF model

An antibody depletion model of CD11b$^+$ myeloid cell under the APAP-induction was used to assess the therapeutic efficacy of CD11b$^+$ myeloid cells. 5 mg/kg of CD11b$^+$ myeloid cell depletion antibody (BioXCell, BE0007) and IgG antibody (BioXCell, BE0090) were intraperitoneally injected 16 h before 500 mg/kg dose of APAP induction. 4 × 10$^9$ EVs were intravenously injected 8 h after APAP induction. To test its depletion for pre-test, liver and spleen tissues were harvested 24 h after APAP induction and dissociated into the single-cell populations utilizing the gentleMACS™ Octo Dissociator with Heaters. The isolated single cells were further stained with a flow cytometric antibody panel. To determine the therapeutic efficacy in the reduction of CD11b$^{+}$ myeloid cells, serum and liver tissues were harvested at 48 h post-APAP induction and analyzed biochemistry (ALT), tissue staining (CD47 IHC) following the same protocol in the method section of manuscript.

**Hemagglutination assay**

For confirming RBC aggregation by EVs or anti-CD47 antibody (Santa Cruz Biotechnology, sc-21786), washed human RBCs (Innovative Research, IWB3ALS40ML) were incubated with Alsever's solution (Sigma-Aldrich, A3551) and centrifuged at 600 g for 5 min at 4 °C, twice. The pellets were then resuspended in PBS and centrifuged again at 600 g for 5 min at 4 °C. Subsequently, the pellets were prepared in a 5 % RBC/PBS solution. The samples were incubated with the indicated quantities of samples for 4 h at 37 °C in U-shaped bottom 96 well microplate. The aggregations were analyzed by flow cytometry or microscopy (Nikon, ECLIPSE Ts2).
